# Molecular mechanisms underlying metamorphosis, including the formation of the prepupa, in the black soldier fly (*Hermetia illucens*)

**DOI:** 10.64898/2026.05.31.729150

**Authors:** Lingtao Zhang, Masami Shimoda, Chieka Minakuchi

## Abstract

The black soldier fly (*Hermetia illucens*, Diptera) undergoes an atypical metamorphic program in which a distinct non-feeding prepupal instar precedes pupation, but its developmental and molecular basis remains poorly understood. Here, we investigated this unusual metamorphic program through integrated developmental and RNAi-based analyses. Postembryonic staging confirmed that the 6^th^ instar feeding larva molts into a non-feeding 7^th^ instar prepupa, whose hardened cuticle subsequently serves as the puparium during intra-puparial development. Expression profiles of four key metamorphic genes revealed a stage-specific *Chinmo*/*Kr-h1*– *Br-c*–*E93* regulatory shift corresponding to larval, prepupal, and pupal/adult development, with sustained *Br-c* expression defining the 7^th^ instar prepupal stage. RNAi-mediated knockdown of *Kr-h1* and/or *Chinmo* induced precocious larval–prepupal metamorphosis, supporting their roles in larval stage maintenance, whereas depletion of *Br-c* or *E93* disrupted prepupal–pupal transition and adult differentiation, respectively, consistent with their functions as pupal and adult specifiers. These results support generally conserved functions of the metamorphic gene network while indicating the absence of a repressive effect of Br-c on *E93* in *H. illucens* prepupae. Together, these findings establish the 7^th^ instar prepupa as an independently regulated transitional stage, providing insight into how metamorphic programs are reorganized to diversify life-history strategies in holometabolous insects.

## Introduction

Insects are the most diverse group of animals, representing over half of all described species on Earth (Belles, 2020). The evolutionary success of insects is closely linked to the key innovations of wings and the pupal stage, leading to the most widely recognized classification of insect metamorphosis into three major types: ametabolan, hemimetabolan, and holometabolan (Truman, 2019). The holometabolous life cycle, which creates an ecological division by separating the juvenile and adult life stages through a non-feeding pupal stage, significantly lowers intraspecific competition and thus facilitates differentiation in ecological niches (Rainford et al., 2014).

From an endocrinological perspective, insect metamorphosis is mainly regulated by two humoral factors: the juvenile hormone (JH) and ecdysteroids (20-hydroxyecdysone (20E), the active form), as well as the downstream metamorphic gene network (MGN) (Truman and Riddiford, 2026). In holometabolous insects, four key transcription factor-encoding genes of the MGN are closely related to the larval–pupal–adult life history: *Krüppel homolog 1* (*Kr-h1*), *Chronologically inappropriate morphogenesis* (*Chinmo*), *Broad-complex* (*Br-c*; also known as *broad*), and *Ecdysone-induced protein 93F* (*E93*). In this scenario, *Kr-h1* acts downstream of the JH receptor complex of Methoprene-tolerant (Met) and Taiman (Tai), thereby mediating the antimetamorphic effect of JH and maintaining the larval status (Minakuchi et al., 2009; Jindra et al., 2015). Recent studies identified *Chinmo* as the larval specifier, promoting larval development from early developmental stages and also maintaining the larval status (Truman and Riddiford, 2022; Chafino et al., 2023; Chen et al., 2024). *Br-c* functions as the pupal specifier, showing upregulated expression during the last larval stage, directing the larval–pupal transition (Karim et al., 1993; Zhou et al., 1998; Zhou and Riddiford, 2002; Konopova and Jindra, 2008). *E93* acts as the adult specifier, which promotes adult differentiation and terminates earlier metamorphic programs (Ureña et al., 2014; Lam et al., 2022; Cruz et al., 2024). The interactive relationships among the key metamorphic genes have been widely studied, establishing and extending the Met–Kr-h1–E93 (MEKRE93) pathway, together constructing a transcriptional regulatory framework that governs stage-specific transitions during insect metamorphosis (Belles and Santos, 2014; Martín et al., 2021; Escudero et al., 2025).

The black soldier fly (*Hermetia illucens*, Diptera: Stratiomyidae) is gathering increased attention nowadays for its broad potential in waste management and industrial applications. Native to the Americas but now widely distributed worldwide (Kaya et al., 2021), *H. illucens* is widely used for the bioconversion of organic waste, as its larvae consume a wide range of substrates and have the potential to provide valuable biomass resources (Nguyen et al., 2015; Surendra et al., 2020). *H. illucens* biomass is rich in protein, fat and minerals, with reports showing 30 ∼ 52% crude protein and 21 ∼ 40% fat contents of dry matter, respectively (Spranghers et al., 2017; Surendra et al., 2020). Utilizing *H. illucens* in biowaste treatment is also environmentally favorable, as its global warming potential (GWP) has been reported to be approximately half that of composting (Mertenat et al., 2019). Due to the above benefits, *H. illucens* is increasingly being applied and accepted, especially as aquafeed or poultry feed (Barragan-Fonseca et al., 2017; Mohan et al., 2022; Oktay et al., 2024). Recent studies have also facilitated the application of genetic tools in this species. RNAi-mediated knockdown has been applied to genes encoding an acyl carrier protein and amino acid transporters (Peng et al., 2023; Liu et al., 2024), and genome-editing approaches have generated productivity-related genetically modified strains, including the wingless and *Higiant* knockout strains (Kou et al., 2024; Wang et al., 2025).

Despite the practical importance and advances in genetic research in *H. illucens*, the developmental and molecular basis of its metamorphosis remains poorly understood. The developmental program in *H. illucens* differs from that of typical holometabolous insects, marked by a molt from the last feeding larval instar into a morphologically and behaviorally distinct instar, which is generally characterized by cuticle sclerotization and darkening, a profound change in the head morphology, the cessation of feeding, and the onset of wandering behavior, with its cuticle later serving as the puparium (Sheppard et al., 1994; Tomberlin et al., 2009; Fabian et al., 2025). Although previous studies have described feeding larval instar identification and intra-puparial development in this species (Kim et al., 2010; Barros-Cordeiro et al., 2014; Ohhara et al., 2026), there is no consensus on the recognition of the instar immediately preceding pupation, as it has been variously referred to as the post-feeding larva (Holmes et al., 2012; Bruno et al., 2020), the final instar larva (Gligorescu et al., 2019; Fabian et al., 2025), or the prepupal stage (Sheppard et al., 1994; Tomberlin et al., 2009; Surendra et al., 2020; Sangkaew et al., 2025), sometimes even misinterpreted for the actual pupa (Kim et al., 2010; Barros et al., 2019). Therefore, the identity of the “prepupal instar” and its role in *H. illucens* metamorphosis are not well characterized, with uncertainty as to whether it constitutes an extension of larval status or marks early acquisition of pupal characteristics. Moreover, it remains largely unexplored how key metamorphic genes of the MGN exert stage-specific effects to regulate the unique metamorphosis program in this species.

In this study, we conducted an in-depth investigation into *H. illucens* metamorphosis through an integrated analysis combining morphology, developmental expression profiling, and RNAi-based functional analyses. Morphological records across postembryonic development confirmed the formation of the puparium via a 6^th^ to 7^th^ instar larval–prepupal molt prior to the onset of wandering behavior, supporting the prepupal instar as a non-feeding, transitional stage that prepares for intra-puparial development. Developmental expression profiling further uncovered a clear stage-specific regulatory shift of *Chinmo*/*Kr-h1*–*Br-c*–*E93* in the metamorphic gene network corresponding to the larval–prepupal–pupal/adult metamorphic program, with elevated *Br-c* expression as a defining molecular feature of the *H. illucens* prepupal stage. Furthermore, RNAi-mediated knockdown experiments confirmed the generally conserved roles of key metamorphic genes in *H. illucens* and suggested a lack of repressive effect of Br-c on *E93* in *H. illucens* prepupae. Together, these findings identify the 7^th^ instar prepupa as a distinct, independently regulated transitional stage in *H. illucens* development, providing insight into how the conserved metamorphic gene network can be reorganized to generate divergent life-history strategies in holometabolous insects.

## Results

### Puparium formation occurs before wandering behavior in *H. illucens*

To characterize the progression of metamorphosis, morphological and behavioral changes were examined throughout postembryonic development in *H. illucens* (Figure 1A). The larval stage of *H. illucens* comprises 6 instars (L1 to L6), during which individuals exhibit continuous feeding behavior. Measurements confirmed that the head capsule width increases progressively across successive instars during the larval stage, consistent with previous reports (Kim et al., 2010), providing a reliable trait for *H. illucens* larval staging (Figure 1—figure supplement 1).

**Figure 1.**
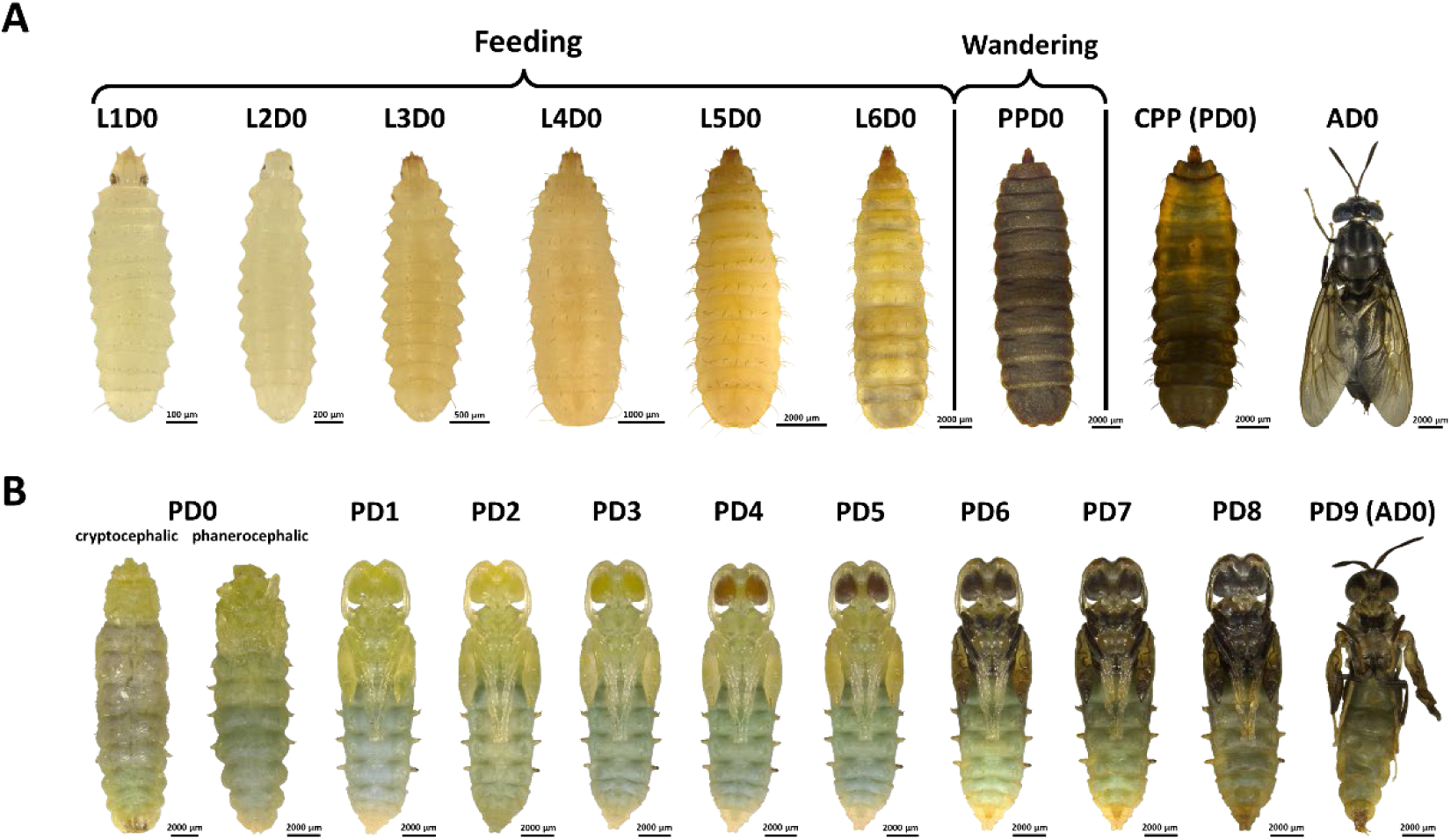
Postembryonic development of *H. illucens*. (**A**) The overall program of postembryonic development, including the feeding larval stage (L1D0 to L6D0), the non-feeding, wandering 7^th^ instar prepupal stage (PPD0), the pupal stage (PD0), and the adult (AD0). (**B**) Intra-puparial development of *H. illucens*, including the cryptocephalic pupa (early PD0), the phanerocephalic pupa (mid-late PD0), the pharate adult (PD1 to PD8), and the newly emerged adult (from PD9 to AD0). L1D0 to PPD0, and AD0 are presented in dorsal view, and the other photos are presented in ventral view. The same individual is exhibited from PD1 to PD9 (AD0).

Following the molt of the 6^th^ instar larva, *H. illucens* enters a distinct 7^th^ prepupal instar (PP) characterized by remarkable morphological changes, cessation of feeding, and the onset of wandering behavior (Figure 1A). The cuticle becomes darkened and hardened at this prepupal stage. A marked reduction in head capsule width was also observed (Figure 1—figure supplement 1), accompanied by pronounced morphological changes in the head structures, including simplified mandibulo-maxillary complex and enlarged eyes (Figure 1—figure supplement 2). The non-feeding prepupae actively exit the feeding substrate and migrate in search of a suitable pupation site.

After wandering and burrowing into the pupation substrate, prepupae gradually became immobile, marking the onset of prepupal–pupal transition. The hard prepupal cuticle serves as the puparium during pupal development. Individuals exhibiting immobility and a transparent dorsal vessel were identified as the cryptocephalic pupae (CPP; early PD0). Subsequent intra-puparial development was documented by removal of the puparium, revealing the transition into the phanerocephalic pupae (mid-late PD0) followed by progressive differentiation into pharate adults (PD1 to PD9). As intra-puparial development gradually proceeds, a yellow-amber-brown change in eye pigmentation was recorded, followed by the onset of body pigmentation (Figure 1B, Supplementary Table S1). These observations demonstrate that in *H. illucens*, puparium formation occurs prior to the onset of wandering behavior, defining a distinct 7^th^ instar non-feeding prepupal stage established through a larval–prepupal molt preceding pupation and intra-puparial development.

### *H. illucens* exhibits unique metamorphic gene network expression patterns during metamorphosis

To elucidate how this unique prepupal-instar-forming life cycle is molecularly regulated, developmental expression profiles of major metamorphosis-associated genes were constructed across the entire postembryonic developmental period, with *Actin-5C* as a reference gene. Alignment of the expression profiles of four key metamorphic genes composing the metamorphic gene network—*Kr-h1*, *Chinmo*, *Br-c*, and *E93*—revealed a clear stage-specific shift in dominant mRNA expression from *Chinmo* and *Kr-h1* during the larval stage to *Br-c* in the prepupal stage, followed by *E93* in pupal and adult stages (Figure 2A).

**Figure 2.**
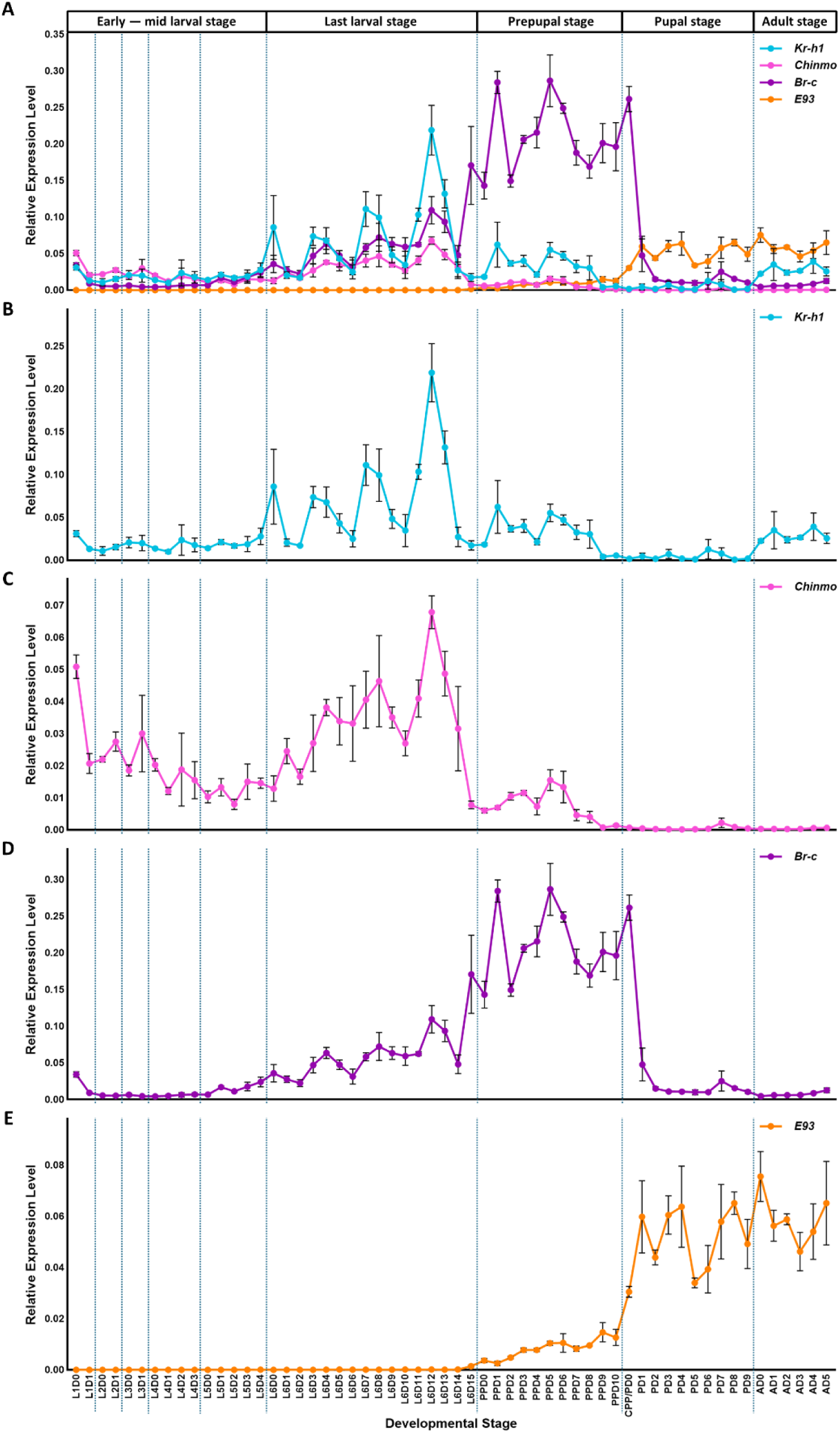
Expression profiles of key metamorphic genes in *H. illucens*. (**A**) Aligned expression profile of genes encoding four key metamorphic genes in *H. illucens*: *Kr-h1*, *Chinmo*, *Br-c* and *E93*. (**B-E**) Separated expression profiles of (**B**) *Kr-h1*, (**C**) *Chinmo*, (**D**) *Br-c*, and (**E**) *E93*. Relative expression level was calculated using the 2^−ΔΔCt^ method, with *Actin-5C* as the reference gene. All data are shown in mean ± SE (n = 3). **Source data 1.** Numerical values for expression profiles shown in Figure 2.

*Kr-h1* expression was at low basal levels during the early-mid larval stage (L1 to L5), showed pronounced pulsatile peaks in the last larval stage (L6), decreased to modest levels upon prepupal entry (PP), declined to near-baseline levels by the late prepupal stage (PPD9), and was re-induced after adult emergence (Figure 2B). *Chinmo* expression was relatively high during the early larval stages (L1 to L3), exhibited a coordinated pattern with *Kr-h1* across the last larval and prepupal stages (L6 to PP), but was not re-induced in adults, indicating a more restricted role of *Chinmo* as the larval specifier (Figure 2C).

*Br-c* displayed a continuously high expression level throughout the entire prepupal stage, and declined sharply upon pupal entry, indicating that elevated *Br-c* expression is a defining molecular feature of the *H. illucens* prepupal stage (Figure 2D). *E93* expression was initiated but remained low during the prepupal stage, but was rapidly upregulated after pupal entry, consistent with a close association with adult differentiation (Figure 2E).

In parallel, developmental expression profiles of the JH/20E receptor-encoding genes *Met* and *EcR*, together with JH/20E degradation enzyme-encoding genes *JHE* and *Cyp18a1*, exhibited additional stage-dependent dynamics, including a prominent *JHE* peak at the end of the prepupal stage and a marked *EcR* peak at the end of the pupal stage, suggesting the elimination of JH signaling before pupation and a high requirement for 20E before adult emergence (Figure 2—figure supplement 1, Figure 2—figure supplement 2).

Together, these results reveal a distinct mRNA expression pattern of the metamorphic gene network during *H. illucens* metamorphosis, characterized by a *Chinmo*/*Kr-h1***–***Br-c***–***E93* regulatory shift corresponding to larval–prepupal–pupal/adult stages, in which the prolonged elevation of *Br-c* expression defines the 7^th^ instar prepupal stage as a unique and independently regulated phase of metamorphosis.

### Knockdown of *Kr-h1* or *Chinmo* induces precocious prepupal entry in *H. illucens*

To examine the functional roles of these stage-specific regulatory factors, RNAi-mediated knockdown experiments were conducted in *H. illucens*. L4D0 larvae were subjected to ds*Kr-h1* or ds*Chinmo* treatment to test the roles of *Kr-h1* and *Chinmo* in maintaining larval identity.

To validate RNAi efficiency, individuals were collected at L4D2 following dsRNA injection, and relative mRNA expression levels of target genes were quantified. Although no clear dose-dependent reduction in transcript levels was observed across the tested concentrations, significant decreases in mRNA expression were detected in the 1500 ng and 3000 ng treatment groups compared with the control group injected with 3000 ng ds*malE* (Figure 3—figure supplement 1). Based on these results, a dose of 2000 ng dsRNA was selected as an appropriate and effective concentration for subsequent functional analyses.

**Figure 3.**
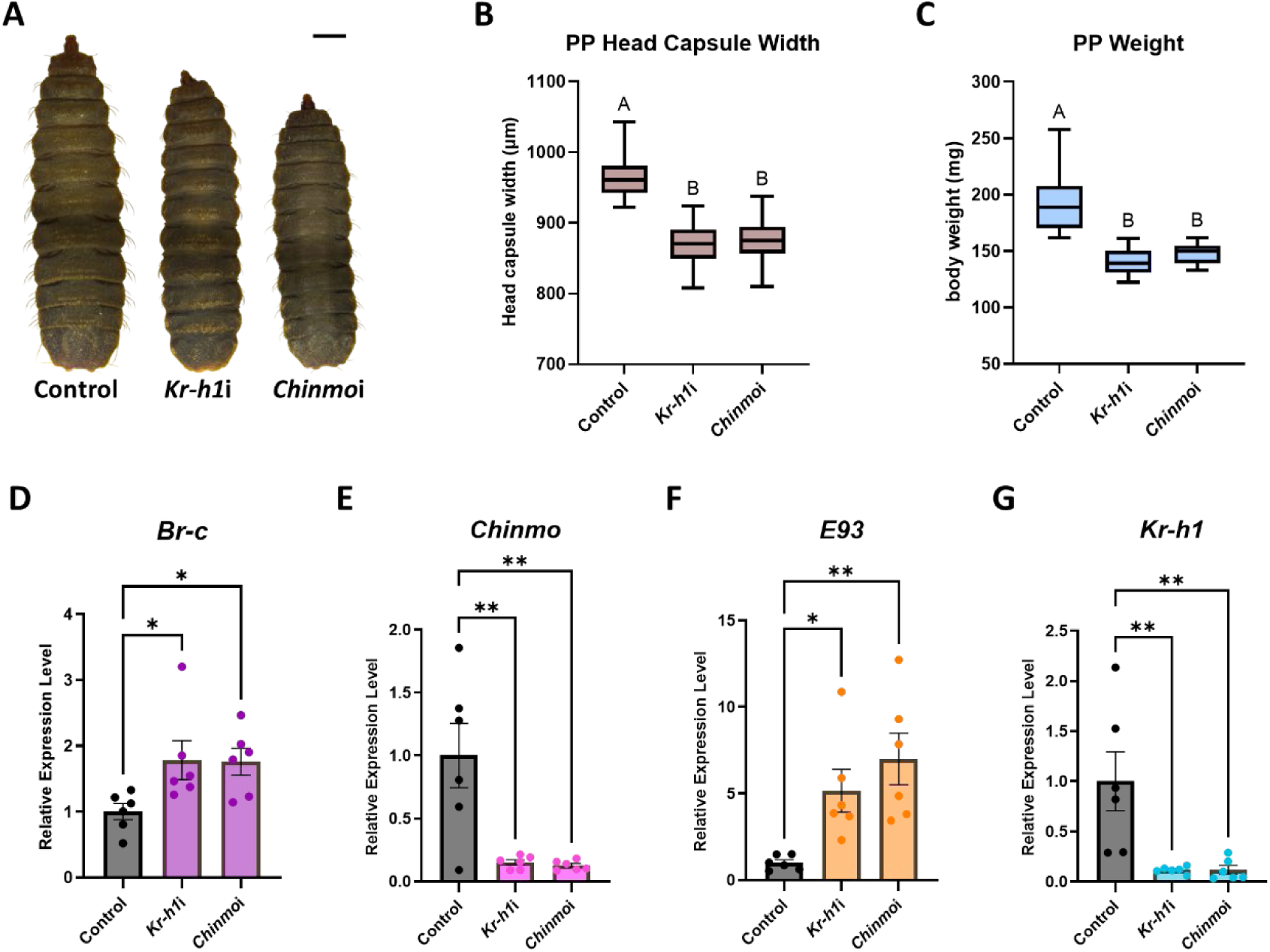
Knockdown of *Kr-h1* or *Chinmo* triggers precocious metamorphosis in *H. illucens*. (**A**) Dorsal view of the control (ds*malE* treated) prepupa, the *Kr-h1*i precocious prepupa, and the *Chinmo*i precocious prepupa. (**B**) Head capsule width comparison of the control (ds*malE* treated) prepupa (n = 58), the *Kr-h1*i precocious prepupa (n = 25), and the *Chinmo*i precocious prepupa (n = 34). (**C**) Body weight comparison of the control (ds*malE* treated) prepupa (n = 58), the *Kr-h1*i precocious prepupa (n = 25), and the *Chinmo*i precocious prepupa (n = 34). (**D**-**G**) Fold changes of relative expression levels in key metamorphic genes: *Br-c* (**D**), *Chinmo* (**E**), *E93* (**F**) and *Kr-h1* (**G**) measured 14 days after dsRNA treatment (mean ± SE, n = 6 for each group). The average relative expression level of the control group was normalized to 1 for analysis. Scale bar: 2000 μm. One-way ANOVA with Tukey’s test was performed for (**B**) and (**C**); groups labeled with different letters are significantly different (*p* < 0.05). One-way ANOVA followed by Dunnett’s multiple comparisons test against the control group was performed for (**D**) to (**G**); Asterisks denote statistical significance: p < 0.05 (*), and p < 0.01 (**).

RNAi-mediated depletion of either *Kr-h1* or *Chinmo* in L4D0 larvae induced precocious prepupal entry after L5, yielding 6^th^ instar prepupae (hereafter referred to as *Kr-h1*i or *Chinmo*i prepupae) instead of the normal 7^th^ instar prepupae (Figure 3A). The proportion of precocious prepupal entry was higher in ds*Chinmo*-treated individuals (68.0%) than in ds*Kr-h1*-treated ones (48.1%), and both treatments also caused a small proportion of L5-to-PP molt failure (Supplementary Table S2). The resulting *Kr-h1*i and *Chinmo*i prepupae exhibited significantly smaller head capsule width and lower body weight compared with control prepupae (Figure 3B & 3C, Supplementary Table S2), indicating premature metamorphic transition prior to completion of ordinary larval growth. However, all *Kr-h1*i and *Chinmo*i prepupae successfully pupated and became adults.

After 14 days following dsRNA injection, 5^th^ instar larvae were collected for mRNA expression level analysis. qRT-PCR further confirmed that in both *Kr-h1*i and *Chinmo*i individuals, the relative expression levels of both *Kr-h1* and *Chinmo* were significantly reduced, whereas *Br-c* and *E93* were significantly upregulated compared with controls (Figure 3D-G), suggesting the repressive role of *Kr-h1* and *Chinmo* to *Br-c* and *E93* during the larval stage. Altogether, these findings demonstrate that *Kr-h1* and *Chinmo* are both required for larval maintenance in *H. illucens*, likely by preventing precocious activation of the prepupal- and adult-associated key regulatory factors *Br-c* and *E93*.

### Double knockdown reveals the cooperative role of *Kr-h1* and *Chinmo* in larval stage maintenance

To further examine whether *Kr-h1* and *Chinmo* act cooperatively in larval stage maintenance, simultaneous knockdown of both genes was performed. Co-injection of ds*Kr-h1* and ds*Chinmo* into L4D0 larvae induced precocious prepupal entry, with 60.0% of treated individuals entering the prepupal stage after L5 (hereafter referred to as *Kr-Chm*i prepupae) (Figure 4A), which was accompanied by a slightly elevated L5-to-PP molt failure rate of 9.1% (Supplementary Table S3). *Kr-Chm*i prepupae also exhibited significantly lower body weight and smaller head capsule width than those of the 4000 ng ds*malE*-treated control group individuals (Figure 4B & 4C, Supplementary Table S3).

**Figure 4.**
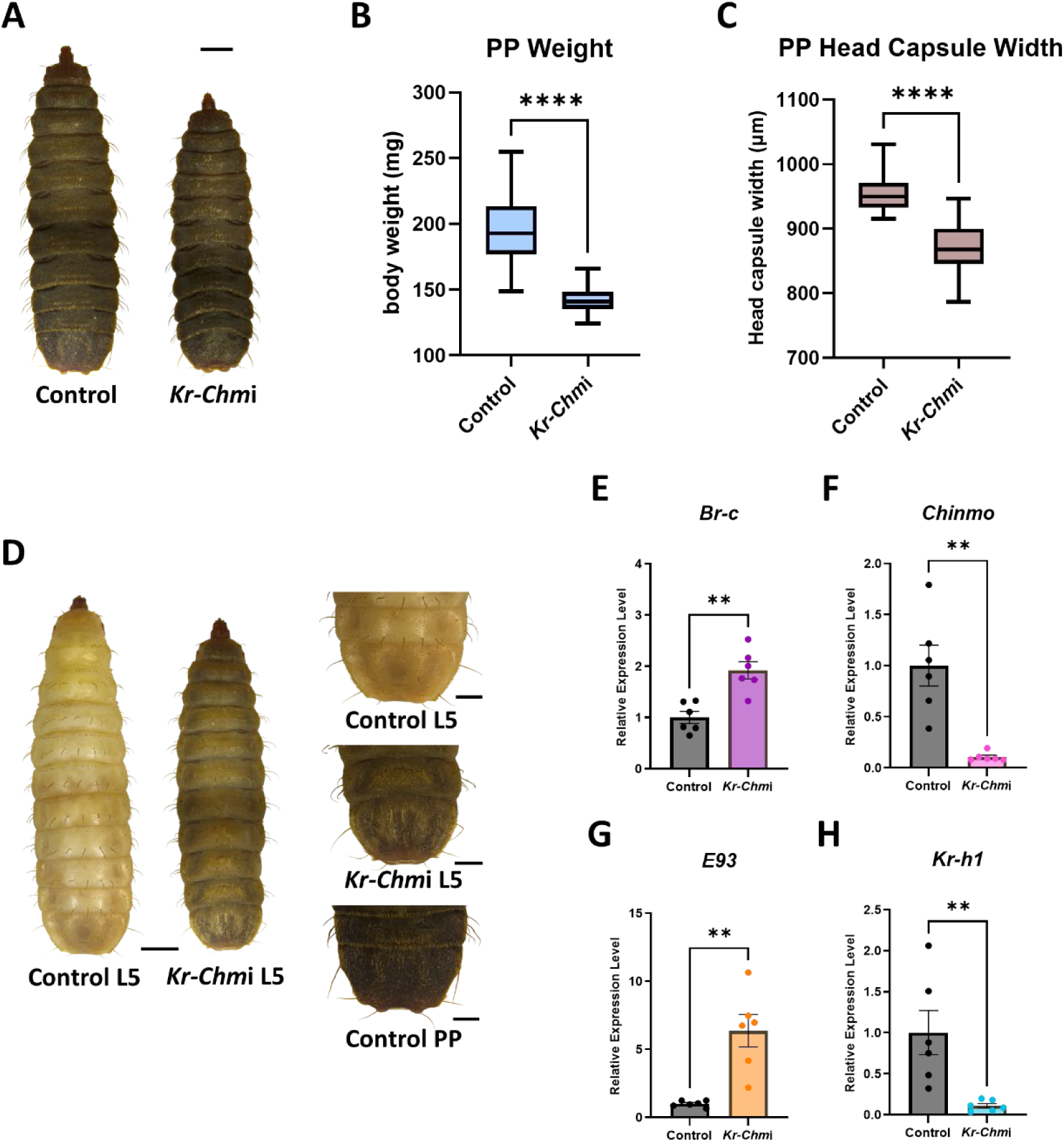
Double knockdown of *Kr-h1* and *Chinmo* triggers precocious metamorphosis and results in PP-like L5 larvae in *H. illucens*. (**A**) Dorsal view of the control (ds*malE* treated) prepupa and the *Kr-Chm*i precocious prepupa. (**B**) Body weight comparison of the control (ds*malE* treated) prepupa (n = 42) and the *Kr-Chm*i precocious prepupa (n = 33). (**C**) Head capsule width comparison of the control (ds*malE* treated) prepupa (n = 42) and the *Kr-Chm*i precocious prepupa (n = 33). (**D**) Morphological comparisons among the prepupal-like L5 larva, the control L5 larva and the control prepupa. Dorsal view of the posterior abdominal segments reveals prepupal-like epidermal phenotype in *Kr-Chm*i L5 larva. (**E**-**H**) Fold changes of relative expression levels in key metamorphic genes: *Br-c* (**E**), *Chinmo* (**F**), *E93* (**G**) and *Kr-h1* (**H**) measured 14 days after dsRNA treatment (mean ± SE, n = 6 for each group). The average relative expression level of the control group was normalized to 1 for analysis. Scale bars: 2000 μm for the whole body, and 1000 μm for body parts. An unpaired Student’s t-test was used to compare the two groups in (**B**), (**C**), (**E**-**H**). Asterisks denote statistical significance: p < 0.05 (*), p < 0.01 (**), p < 0.001 (***), and p < 0.0001 (****).

Intriguingly, 21.2% of the *Kr-Chm*i individuals (7 of 33) exhibited a distinct prepupal-like epidermal phenotype at L5, which was observed exclusively in the double-knockdown group, characterized by darkened body coloration and excessive abdominal setae (Figure 4D), indicating a stronger disruption of larval epidermal identity compared with single knockdown treatments. qRT-PCR analysis conducted using L5 larvae at 14 days after dsRNA injection also revealed reduced expression of *Kr-h1* and *Chinmo*, accompanied by significant upregulation of *Br-c* and *E93*, with a slightly more stable elevation of *Br-c* relative to single knockdowns (Figure 4E-H). These results indicate that *Kr-h1* and *Chinmo* cooperatively maintain the larval status in *H. illucens*, and that their combined depletion promotes premature metamorphic progression while more strongly impairing larval identity.

### *Br-c* is essential for prepupal-pupal transition in *H. illucens*

Given the markedly elevated expression of *Br-c* from the beginning of the prepupal stage, RNAi-mediated knockdown was conducted in PPD0 individuals to assess the role of Br-c during *H. illucens* prepupal development. RNAi efficiency in prepupae was confirmed by qRT-PCR at PPD2, showing that *Br-c* transcript levels were significantly reduced in ds*Br-c*-treated individuals to approximately 0.65-fold of control levels (Figure 5—figure supplement 1).

**Figure 5.**
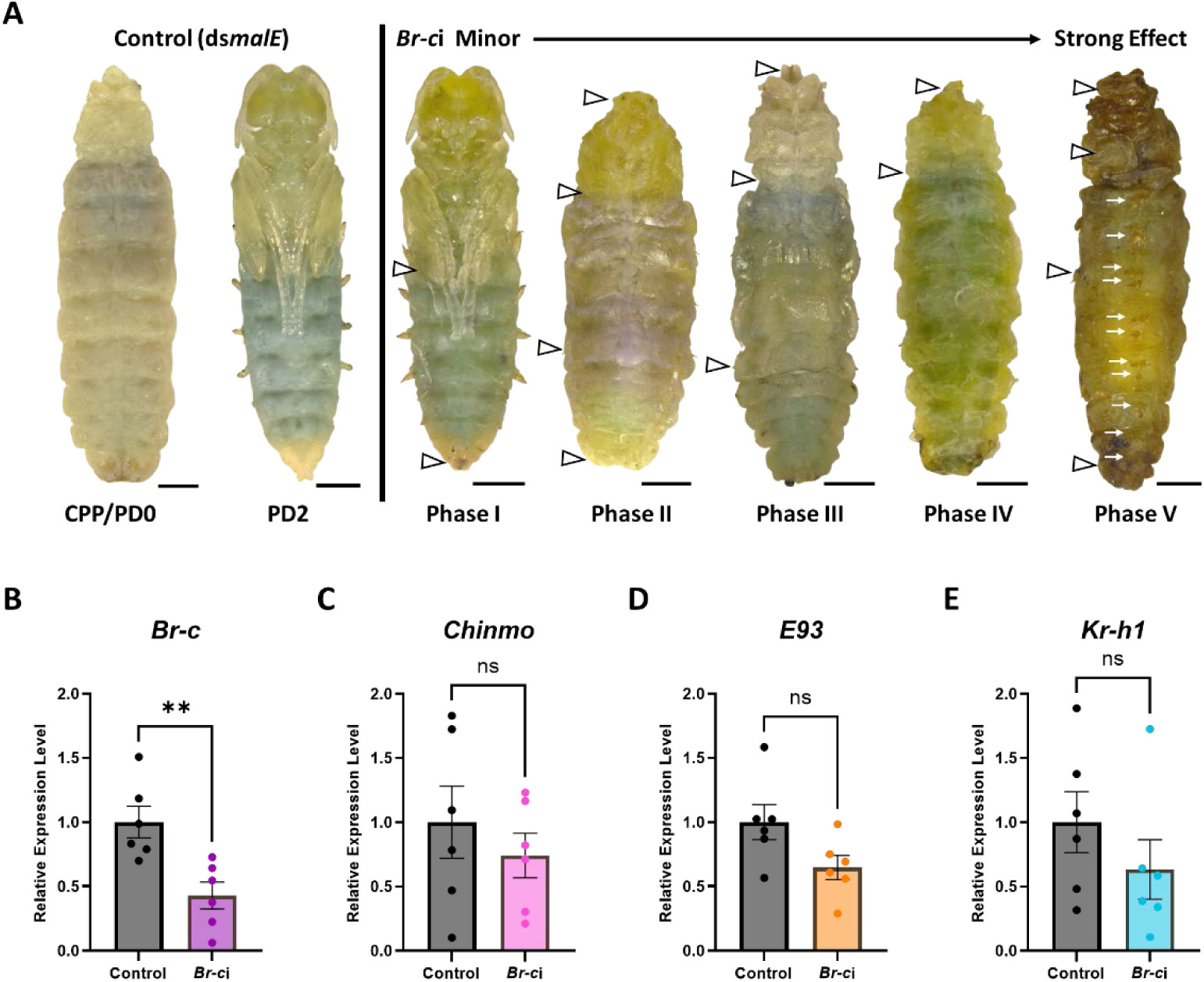
Knockdown of *Br-c* impedes prepupal-pupal metamorphosis in *H. illucens*. (**A**) Phenotypes obtained from ds*malE*-injected control individuals and ds*Br-c*-injected RNAi individuals (*Br-c*i): cryptocephalic pupa (CPP/PD0), pharate adult (PD2), and *Br-c*i phases I-V individuals. Phase I: pharate adult wings were expanded but flat, attaching to the 1^st^ and 2^nd^ abdominal segments, and terminal abdominal projections were incompletely developed; Phase II: overall development was arrested at phanerocephalic pupal stage. Eyes were incompletely developed, legs were formed but unextended, wings were partially expanded but flat, attaching to the metathorax and the 1^st^ abdominal segment. Abdominal respiratory tubes were incompletely formed, and the abdominal terminal segment was flat and larval-like. Phase III: overall development was arrested at the cryptocephalic pupal stage. Head and legs were differentiated but undeveloped, wings were partially expanded but small and flat, attaching to the metathorax. Abdominal respiratory tubes were unformed. Phase IV: less differentiated than phase III, the head and legs were almost undifferentiated, and the wing buds were unexpanded. Phase V: overall development was arrested at the phanerocephalic pupal stage, similar to phase II, except that wings failed to reach the 1^st^ abdominal segment. Prepupal-like setae appeared on each abdominal segment. (**B**-**E**) Fold changes of relative expression levels in key metamorphic genes: *Br-c* (**B**), *Chinmo* (**C**), *E93* (**D**) and *Kr-h1* (**E**) measured 7 days after dsRNA treatment (mean ± SE, n = 6 for each group). The average relative expression level of the control group was normalized to 1 for analysis. Important morphological characteristics described are labeled with white arrowheads, and prepupal-like setae are indicated by white arrows. Scale bars: 2000 μm. An unpaired Student’s t-test was used to compare the two groups in (**B**-**E**). Asterisks denote statistical significance: p < 0.05 (*), p < 0.01 (**), and p < 0.001 (***).

Notably, whereas all control individuals developed normally and completed adult emergence, all ds*Br-c*-treated (*Br-c*i) individuals exhibited developmental arrest and failed to form normal pharate adults. The arrested *Br-c*i individuals were classified into five phenotypic phases according to the degree of developmental progression, with phase I corresponding to the least severe and phase V to the most severe phenotype (Figure 5A, Supplementary Table S4). *Br-c*i phase I individuals reached a pharate adult-like condition but exhibited flattened wings attached to the abdomen and incompletely developed terminal abdominal projections. Phase II and phase III individuals were arrested at the phanerocephalic and cryptocephalic pupal stages, respectively, while phase IV individuals were arrested at the earliest developmental stage, with poorly differentiated head, legs, and wing buds (Figure 5A).

The most severe *Br-c*i phase V phenotype, although progressing to a morphology corresponding to the phanerocephalic pupal stage, was distinguished by the appearance of prepupal-like setae on abdominal segments (Figure 5A, Figure 5—figure supplement 2). This retention of prepupal epidermal characteristics despite pupal progression suggests disruption of the normal prepupal-pupal transition under the depletion of *Br-c*. Additional higher-dose (5000 ng) and repeated ds*Br-c* injection (2000 ng × 3) treatments also produced complete developmental arrest in all ds*Br-c*-treated individuals, with a higher proportion of phase III–V phenotypes than in the standard 2000 ng treatment (Supplementary Table S4).

qRT-PCR analysis performed in prepupae 7 days after dsRNA injection confirmed that *Br-c* expression levels were significantly reduced in *Br-c*i individuals compared with ds*malE*-treated controls (Figure 5B). However, the expression levels of *Chinmo*, *E93*, and *Kr-h1* did not differ significantly between the two groups (Figure 5C–E), indicating that the role of *Br-c* in *H. illucens* prepupal–pupal transition is more likely through the direct regulation of downstream pupal differentiation programs, rather than repressing other key components of the MGN. These results support a specific requirement for *Br-c* in directing the metamorphic transition in *H. illucens* prepupa.

### *E93* is essential for adult differentiation in *H. illucens*

Because the exact segmental position is difficult to determine externally across the puparium in *H. illucens*, ds*E93* was injected into late prepupae at PPD7 to examine the role of E93 during the pupal–adult transition. *E93* transcript levels were significantly reduced in ds*E93*-treated individuals to ∼0.64-fold of control levels, showing valid RNAi efficiency (Figure 6—figure supplement 1).

**Figure 6.**
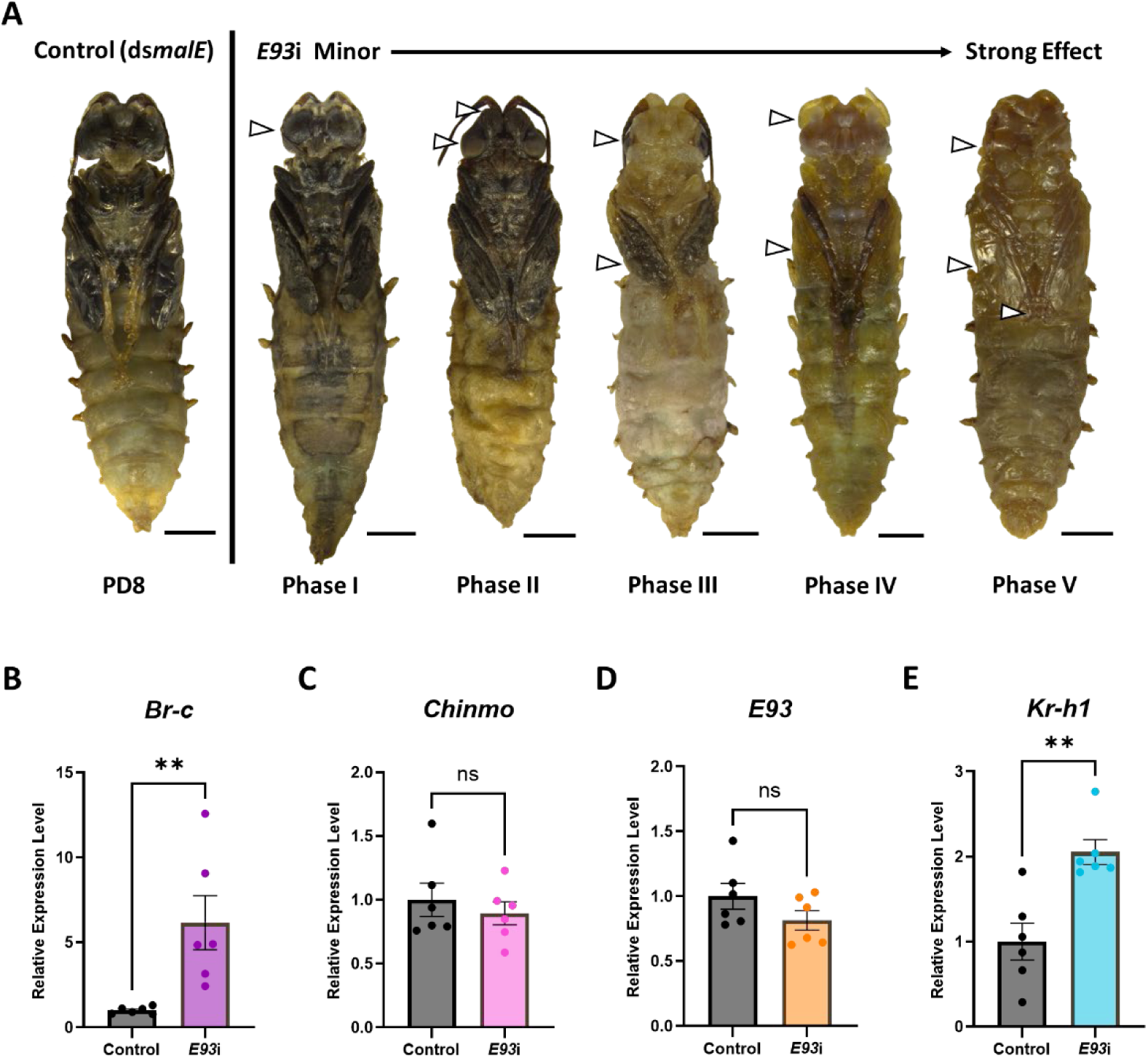
Knockdown of *E93* inhibits adult formation in *H. illucens*. (**A**) Phenotypes obtained from ds*malE*-injected control individuals (pharate adult at PD8) and ds*E93*-injected RNAi individuals (*E93*i phases I-V). Phase I: all tissues normally developed except slightly smaller compound eyes in some individuals. Pupae failed to complete eclosion and were trapped in the puparium. Phase II: all tissues were generally well developed, except that the compound eyes were markedly reduced in size. An ectopic swelling appeared between the antennal bases. Phase III: body pigmentation was restricted to antennae, part of the eyes, and wings. Pigmented range in eyes was significantly small, wings were partially expanded but flat, attaching to the metathorax and the 1^st^ abdominal segment. Phase IV: Body pigmentation was generally absent. Antennae were curved, and wings were partially expanded but flat and transparent. Phase V: Body pigmentation was totally absent. The head was incompletely developed, and the wings were partially expanded but flat and transparent. Legs were incompletely extended, with the posterior end attaching to the apex of the wings. (**B**-**E**) Fold changes of relative expression levels in key metamorphic genes: *Br-c* (**B**), *Chinmo* (**C**), *E93* (**D**) and *Kr-h1* (**E**) measured at PD3 (mean ± SE, n = 6 for each group). Important morphological characteristics described are labeled with white arrowheads. Scale bars: 2000 μm. An unpaired Student’s t-test was used to compare the two groups in (**B**-**E**). Asterisks denote statistical significance: p < 0.05 (*), and p < 0.01 (**).

Under the standard treatment condition (2000 ng dsRNA injection), all ds*E93*-treated individuals succeeded in progression into the pharate adult stage. However, only 35.7% of treated individuals completed adult eclosion, whereas the remaining 64.3% were arrested during the pupal–adult transition (Supplementary Table S5). These arrested individuals (hereafter referred to as *E93*i pupae) were also classified into five phenotypic phases according to the severity of adult differentiation defects (Figure 6A). *E93*i Phase I individuals showed the mildest phenotype and largely resembled control PD8 pharate adults, but failed to emerge and in some cases exhibited slightly reduced compound eyes. Phase II individuals additionally developed a conspicuous ectopic swelling between the antennal bases. In phase III individuals, body pigmentation was restricted to the antennae, parts of the compound eyes, and the partially expanded wings, which remained flat against the metathorax and first abdominal segment. Phase IV individuals exhibited more severe defects, including loss of most body pigmentation, curved antennae, and transparent incompletely expanded wings. The most severely affected phase V individuals exhibited nearly complete loss of pigmentation, incomplete head development, partially expanded but transparent wings, and incompletely extended legs with their posterior ends attached to the wing apex (Figure 6A).

A chronological disorder of tissue development was also observed in phase III *E93*i individuals when puparium removal was performed at PD14. In these individuals, eye pigmentation was confined to the outer margins of the eyes and was amber in color, corresponding to an earlier developmental stage (∼PD3 in controls). In contrast, the wings and parts of the legs already showed deep black pigmentation typical of late pharate adult stages (Figure 6—figure supplement 2). Additional higher-dose (5000 ng) ds*E93* treatment further reduced the normal eclosion rate to 16.7% and shifted the phenotype distribution toward more severe phases III–V, whereas repeated ds*E93* injection (2000 ng × 3 at PPD6, PPD8, and PPD10) did not significantly increase pupal arrest rate (66.7%) (Supplementary Table S5).

qRT-PCR analysis at PD3 revealed that, although no significant differences were detected in *Chinmo* and *E93* transcript levels, the expression levels of *Br-c* and *Kr-h1* were significantly elevated in ds*E93*-treated individuals compared with controls (Figure 6B–E). These results indicate that *E93* exerts a repressive effect on *Br-c* and *Kr-h1* during the pupal–adult transition. Collectively, these results indicate that *E93* is not required for progression to the pharate adult stage, but functions as the adult specifier in *H. illucens* by suppressing *Br-c* and *Kr-h1*, thereby promoting adult differentiation and terminating earlier metamorphic programs.

## Discussion

In this study, we demonstrate that *H. illucens* undergoes a distinct prepupal-instar-forming metamorphosis, in which the puparium is formed through a larval–prepupal molt prior to wandering behavior and the 7^th^ instar prepupa functions as a transitional stage preceding pupation. Functions of the four key metamorphic genes are generally conserved in *H. illucens*, with a sustained high expression level of *Br-c* defining the molecular feature of the prepupal stage, forming a stage-specific *Chinmo/Kr-h1*–*Br-c*–*E93* regulatory shift across postembryonic development. These findings support the view that the 7th instar prepupa is an independently regulated transitional phase in *H. illucens* development, and provide a molecular framework for understanding this unusual metamorphic program.

Wandering behavior is widely observed in holometabolous insects during a non-feeding period prior to pupation, particularly in Diptera and Lepidoptera (Denlinger and Zdarek, 1994; Wiklund et al., 2017). This period is of great ecological significance, as it is widely regarded as an antipredator strategy (Lindstedt et al., 2019). In most holometabolous insects, the post-feeding wandering period occurs at the end of the final larval instar after surpassing the threshold size for metamorphosis (minimal viable weight, MVW), without an intervening molt (Mirth and Riddiford, 2007). However, an important exception is found in sawflies (Hymenoptera), an ancestral lineage of holometabolous insects, in which larvae molt into a distinct non-feeding wandering instar termed “eonymph,” “pronymph,” or “prepupa” (Hara et al., 2023; Suzuki, 2026). In this study, we confirmed that the 6^th^ instar larva of *H. illucens* molts into a non-feeding, wandering 7^th^ instar prepupa. After reaching a suitable pupation site, the prepupa ceases wandering, and the metamorphic transition occurs at the final phase of this stage, with prepupal cuticle serving as the puparium. A postembryonic developmental model was then established for *H. illucens* in comparison with typical dipteran insects represented by *Drosophila melanogaster*, in which a larval–prepupal molt is inserted between the larval stage and pupal formation in *H. illucens*, during which the puparium epidermis is established (Figure 7A). A recent work identified the MVW of *H. illucens* to be 83 mg, overlapping the critical weight, as a threshold size inducing the larval-prepupal (6^th^-7^th^ instar) molt (Ohhara et al., 2026). This further supports the view that the 7^th^ instar prepupal stage represents a key developmental transition in the metamorphosis of *H. illucens*. The ecological functions of the atypical prepupal instar, as well as the evolutionary basis for the emergence (or reacquisition) of this life-history strategy resembling ancestral holometabolous insects, remain unresolved and warrant further investigation.

**Figure 7.**
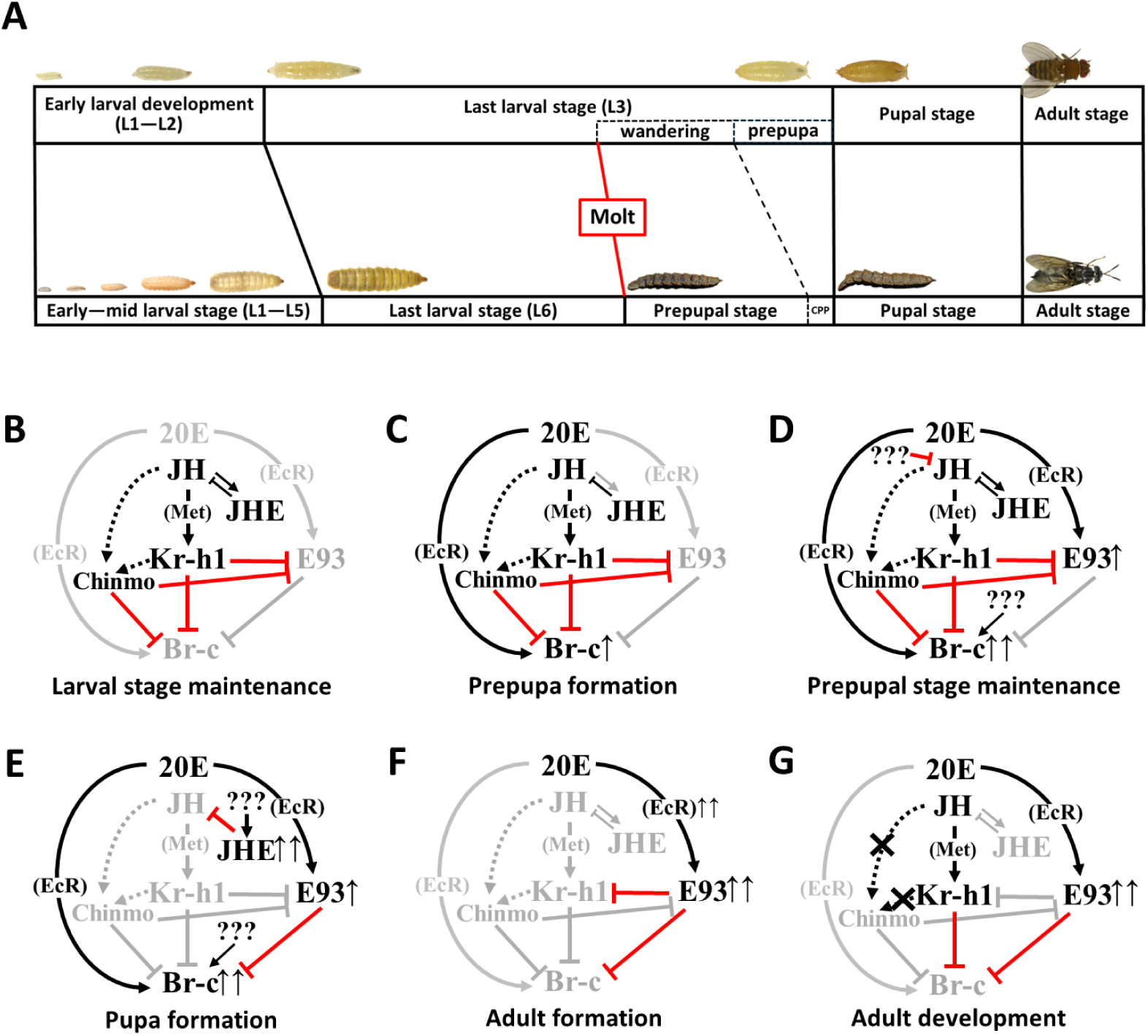
Postembryonic developmental model and schematic diagrams of molecular mechanisms underlying growth and metamorphosis in *Hermetia illucens*. (**A**) The postembryonic developmental model of *H. illucens* in comparison with *D. melanogaster*. Solid lines between stages indicate stage transitions achieved through molting or pupation, whereas dotted lines indicate stage transitions achieved without molting. The model of *D. melanogaster* is adapted from (Martín et al., 2021). (**B-G**) Schematic diagrams of the molecular mechanism underlying growth and metamorphosis in *H. illucens*. (**B**) Larval stage maintenance: a high JH titer in the hemolymph induces the expression of *Kr-h1*, and possibly *Chinmo* as well, as a secondary effect. The Kr-h1 protein suppresses the expression of *Br-c* and *E93*, with Chinmo also suppressing *Br-c*, thereby maintaining the larval status. (**C**) Prepupa formation: 20E signaling induces gradual expression of *Br-c* in the final instar larva. The coexistence of both JH and 20E signaling triggers the larval-prepupal metamorphosis. (**D**) Prepupa maintenance: *Br-c* expression is maintained at a high level throughout the prepupal stage. The JH signal is low but still present, inducing the low-level *Chinmo* and *Kr-h1* expression. (**E**) Pupa formation: The loss of Kr-h1 and Chinmo eliminates the prepupal stage maintenance effect, allowing 20E to induce a gradually increased expression of *E93*. Br-c plays a major role in inducing the prepupal-pupal metamorphosis. (**F**) Adult formation: the elevated presence of E93 protein during the pupal stage represses *Br-c* expression while promoting adult tissue development, triggering pupal-adult metamorphosis. (**G**) Adult development: the JH signal reappears in adults, inducing the upregulation of *Kr-h1*, whereas *Chinmo* expression remains strongly repressed. The coexistence of Kr-h1 and E93 together promotes adult development and maturation.

A sequential expression shift of *Chinmo*/*Kr-h1***–***Br-c***–***E93* was observed corresponding to the larval–prepupal–pupal/adult stage progression in *H. illucens*, with sustained *Br-c* expression emerging as the most distinctive molecular feature of the 7^th^ instar prepupal stage. The expression pattern of *Kr-h1* suggests dynamic changes in JH signaling, with pronounced JH activity during the last larval stage, reduced but persistent activity during early-to-mid prepupal development, a marked reduction upon pupal commitment, and reactivation after adult emergence. This low but persistent prepupal *Kr-h1* expression, together with modest *Chinmo* expression from PPD0 to PPD8, may be correlated to a JH-dependent antimetamorphic effect that maintains the prepupal stage. The expression pattern of *Br-c* in *H. illucens* differs markedly from that of typical holometabolous insects, in which *Br-c* is usually co-expressed with *Kr-h1* as a short temporal surge during the prepupal period (Ureña et al., 2016; Truman and Riddiford, 2026; Suzuki, 2026). In *D. melanogaster*, this stage-specific *Kr-h1* surge is 20E-induced (Pecasse et al., 2000; Minakuchi et al., 2008), and it is further revealed that the 20E-EcR/USP-Met-Tai induces *Kr-h1* expression, and therefore upregulates *Br-c* expression via the positive Kr-h1 binding sites in *Br-c* enhancers during larval–prepupal transition (Long et al., 2025). In *H. illucens*, however, *Br-c* was not co-expressed with *Kr-h1* at either larval-prepupal transition (L6D14–PPD0) or prepupal-pupal transition (PPD9–CPP/PD0), nor was its expression pattern consistent with those of *Met* or *EcR* during prepupal development. Therefore, the prepupal-stage-associated elevation of *Br-c* in *H. illucens* is unlikely to be explained by the canonical 20E-induced *Kr-h1*–dependent regulatory mechanism described in *D. melanogaster*. Given that *Br-c* encodes multiple zinc finger–based splice isoforms (Suzuki et al., 2008), isoform-specific expression dynamics may also help resolve the unexplained prepupal expression pattern. Further elucidation of the involvement of JH signaling during the prepupal stage and the transcriptional regulatory mechanism of *Br-c* may provide critical insight into the functional role of this stage in *H. illucens* development.

It is well documented that the four key members of the MGN mutually exert repressive effects on each other to support metamorphic development. Representative examples in holometabolous insects include the repressive effect of Kr-h1 on *Br-c* and *E93* in *Bombyx mori* cell line (Kayukawa et al., 2016, 2017), that of Chinmo on *Br-c* and *E93* in *Spodoptera frugiperda* early-mid larva and *D. melanogaster*, (Chafino et al., 2023; Truman and Riddiford, 2022; Chen et al., 2024), that of Br-c on *Chinmo* in *D. melanogaster* mid-late L3 wing imaginal disc cells (Narbonne-Reveau and Maurange, 2019), that of Br-c on *E93* in *S. frugiperda* final instar larva (Liu et al., 2025), and that of E93 on *Kr-h1* and *Br-c* in *Tribolium castaneum* and *D. melanogaster* pupae (Ureña et al., 2014). RNAi experiments confirmed that the functions of the key metamorphic genes comprising the MGN are generally conserved in *H. illucens*. Notably, *Kr-h1*i, *Chinmo*i, and *Kr-Chm*i individuals all exhibited simultaneous reduction of *Kr-h1* and *Chinmo* levels, together with upregulation of *Br-c* and *E93*. Since there is currently no evidence supporting *Br-c* or *E93* repressing *Chinmo*, these coordinated changes are more likely to reflect secondary effects of disrupted larval identity maintenance than direct regulatory interactions within the MGN. Nevertheless, the prepupal-like setae phenotype observed exclusively in *Kr-Chm*i individuals supports a cooperative role of *Kr-h1* and *Chinmo* in larval maintenance. In *T. castaneum*, depletion of *Br-c* was reported to induce a “larval–adult” mosaic phenotype (Konopova and Jindra, 2008; Suzuki et al., 2008), and a similar phenotype accompanied by elevated *E93* expression was observed in *S. frugiperda* (Liu et al., 2025). In contrast, neither adult-like morphology nor increased *E93* expression was detected in *H. illucens Br-c*i prepupae. These differences suggest that *Br-c* does not exert a repressive effect on *E93* in *H. illucens* prepupae, highlighting a regulatory mechanism distinct from that described in other holometabolous insects. The repressive effect of E93 on *Kr-h1* and *Br-c* was confirmed by upregulated *Kr-h1* and *Br-c* levels in *E93*i pupae. Notably, a single injection of 5000 ng ds*E93* at PPD7 induced stronger phenotypic effects than repeated injections of 2000 ng ds*E93* at PPD6, PPD8, and PPD10. This observation suggests that *E93* function may be particularly sensitive to depletion within a narrow developmental window, during which *E93* activity is critical for adult differentiation.

Combining the findings in this study, schematic diagrams are constructed to illustrate the molecular mechanism underlying growth and metamorphosis in *H. illucens* (Figure 7B-G). Our results support a model in which *H. illucens* does not simply follow the typical larva–pupa– adult sequence, but inserts a distinct 7^th^ instar prepupal stage between larval growth and pupal morphogenesis. This wandering-specific prepupal stage is molecularly defined by sustained *Br-c* expression together with low but persistent *Kr-h1* and *Chinmo* expression prior to the subsequent rise of *E93*. Functional analyses further demonstrate that *Kr-h1* and *Chinmo* cooperatively maintain larval identity, *Br-c* is necessary for the prepupal–pupal transition, and *E93* is required for adult differentiation. Together, these findings reveal that the prepupal-instar-forming metamorphosis of *H. illucens* is regulated through a distinct stage-specific configuration of the metamorphic gene network, providing a mechanistic basis for understanding how metamorphic programs can be reorganized to diversify life-history strategies in holometabolous insects.

## Materials and Methods

### Insect rearing and staging

Fresh eggs were collected from adults reared in insect cages using a commercial *H. illucens* egg collector and chicken manure as an oviposition attractant. The collector was placed into a large plastic cup (Φ129 × 97 mm) filled with water to a depth of ∼1 cm for egg incubation. Freshly hatched L1D0 larvae were collected from the cup after 3 days of hatching.

For routine rearing, around 400 freshly hatched L1D0 larvae were transferred into a plastic container (111 × 81 × 46 mm) and provided with 60 g of artificial diet for the first 7 days (the composition of the artificial diet is shown in Supplementary Table S6). Two sets of 60 L4D0 (D7) larvae were collected weekly; each set was transferred into a medium plastic cup (Φ101 × 80 mm) and provided with 240 g of artificial diet (4 g/larva) for the next two weeks. L6 (D21) larvae were counted and transferred into another medium plastic cup and given 2 g/larva artificial diet. PPD0 prepupae were collected daily after D21 and transferred into a small plastic cup (Φ101 × 40 mm) filled with 20 g of vermiculite and 20 g of water (50% water content) until eclosion. Newly emerged adults were transferred daily to a new medium plastic cup with a wet paper towel underneath.

For head capsule measurements, 60 freshly hatched L1D0 larvae were transferred into a plastic container (111 × 81 × 46 mm) and provided with 20 g of artificial diet. Molting events were confirmed daily by identifying exuviae in the rearing substrate. Individuals of different instars were separated into multiple containers with 20 g of artificial diet each day. PPD0 prepupae were collected daily since the first appearance and transferred into a small plastic cup (Φ101 × 40 mm) filled with 20 g of vermiculite and 20 g of water (50% water content) until eclosion. Photographs of head morphology were taken using a stereo zoom microscope (Axio Zoom V16; ZEISS) at L1D0 and within 24 h after every molt until PPD0, and measurements of head capsule width were conducted using ZEISS ZEN 3.5 (blue edition). Head capsules of L6D0 and PPD0 individuals were imaged with a tabletop SEM (TM4000Plus II; HITACHI).

The entire rearing procedure was conducted under 27±1℃, L:D = 16:8.

### Pupal entry identification and puparium removal

Individuals of the cryptocephalic pupal stage (CPP) were identified using a combination of mobility test and morphological confirmation. PPD0 prepupae were collected and transferred into small plastic cups (Φ101 × 40 mm) filled with 20 g of vermiculite and 20 g of water (50% water content) for 1 week. The mobility of individuals from PPD7 was confirmed by placing all individuals on the surface of the vermiculite and settling the cup in a dark place for 30 minutes. Immobile individuals were illuminated from the dorsal side to confirm the transparency of the dorsal vessel using an LED flashlight. Individuals showing immobility, unfolded abdominal terminal segments, and transparent dorsal vessels were identified as the CPP (PD0) individuals. All individuals showed folded abdominal terminal segments 1 day after CPP, which were identified as PD1 stage individuals.

Puparia were affixed to double-sided tape on a piece of paper, with the ventral side facing up. A corneal scissor was used to cut open the cavity at the end of the abdomen. Two pairs of tweezers (No. 5; DUMONT) were used to peel away the epidermis while stabilizing the rest of the body. Transparent tape was used to secure the separated epidermis to the paper. After completely separating the ventral puparium epidermis, the pupa was collected using a pair of tweezers with rubber tubes attached to the head.

### Intra-puparial development observation

Puparia were transferred into 6-well tissue culture test plates (TPP^®^) with wet Kimwipes (Nippon Paper Crecia) settled on the bottom of each well for individual staging and observation. 20 individuals from PD1 and 10 individuals from CPP were examined daily, and wet Kimwipes were replaced every day. Photographs were taken using a stereo zoom microscope (Axio Zoom.V16; ZEISS).

### RNA extraction and cDNA synthesis

Staging was conducted daily to randomly collect three biological replicates for RNA extraction. Sex was not determined for larvae, prepupae, and pupae, as external sexing is not possible at these stages. Adults were also collected without separating males and females.

Insect tissue samples were prepared and dissected as needed (described in Supplementary Table S7). Total RNA extraction was conducted using TRIzol Reagent (Invitrogen). RNA product obtained from each sample was dissolved in 40 μL RNase-free H_2_O, quantified with a spectrophotometer (BioSpec-nano; SHIMADZU), and stored at −80℃. cDNA was synthesized from ∼800 ng total RNA using PrimeScript™ RT reagent Kit with gDNA Eraser (TaKaRa) according to the manufacturer’s instructions and stored at −20°C until use.

### qRT-PCR

Homology searches were conducted using *Drosophila melanogaster* gene and amino acid sequences as queries in NCBI BLAST. The resulting homologous sequences from *Hermetia illucens* were used to design primers with Primer3 (https://primer3.ut.ee/), and primer specificity was subsequently checked with Primer-BLAST. Primers were designed in a conserved region common to all annotated transcript isoforms. Primer sequences for qRT-PCR are shown in Supplementary Table S8.

For expression profiles of *Kr-h1*, *Br-c*, *E93*, and *JHE*, TB Green Premix Ex Taq II (TaKaRa) was used for qRT-PCR in Thermal Cycler Dice Real Time System TP800 (TaKaRa) following the manufacturer’s instructions. For other experiments in this study, THUNDERBIRD^®^ Next SYBR™ qPCR Mix (TOYOBO) was used for qRT-PCR in QuantStudio 3 real-time PCR system (ThermoFisher) following the manufacturer’s instructions. The PCR conditions are as follows: 95 ℃ for 30 s (hold stage), 40 cycles of 95 ℃ for 5 s, 60 ℃ for 30 s, and 95 ℃ for 1 s (PCR stage); 95 ℃ for 1 s, 60 ℃ for 20 s, and 95 ℃ for 1 s (melt curve stage). Relative expression levels were calculated using the ΔΔCt method with *Actin-5C* as the reference gene. Analysis was performed for three biological replicates, and the average and standard error were calculated.

### dsRNA synthesis

Double-stranded DNA templates with opposing T7 promoters at the 5’ ends of each strand (5’-TAATACGACTCACTATAGGG-3’) were synthesized for each target gene by PCR using TaKaRa Ex Taq^®^ Hot Start Version (TaKaRa) with cDNA samples and primers listed in Supplementary Table S9. PCR was performed with an initial denaturation at 94 °C for 2 min, followed by 10 cycles of 94 °C for 15 s, 55 °C for 30 s, and 72 °C for 1 min; 30 cycles of 94 °C for 15 s, 60 °C for 30 s, and 72 °C for 1 min; and a final extension at 72 °C for 1 min.

DNA templates obtained were purified using QIAquick PCR Purification Kit (QIAGEN), and then used for dsRNA synthesis with MEGAscript RNAi Kit (Thermo Fisher Scientific). As a negative control, a T7 promoter-tagged DNA template was amplified from the pMAL-c2X plasmid (New England Biolabs) (RRID:Addgene_75286) using primers targeting the gene *maltose-binding protein* of *Escherichia coli* (*malE*), purified, and subjected to dsRNA synthesis in the same manner.

All dsRNA samples obtained were quantified with a spectrophotometer (BioSpec-nano; SHIMADZU), diluted to 3000 ng/μL with nuclease-free water, and stored at −20℃ until injection.

### dsRNA injection

For L4D0 larvae, individuals were randomly collected from the same rearing container, anesthetized on ice for 15 min, and then affixed to a double-sided tape attached to the bottom of a Petri dish (Φ90 × 15 mm; AS ONE). dsRNA solution diluted with nuclease-free water to the desired concentration was loaded into a 10 μL microinjector (Φ0.26 mm; Bolige). Each individual was injected with 2.0 μL of dsRNA solution, and then the Petri dish was placed on ice for 10 min. The dsRNA-injected larvae were then removed from the double-sided tape using a pair of tweezers with rubber tubes attached to the head and placed into a medium plastic cup containing 5 g/larva artificial diet.

For prepupae, dsRNA injection was conducted in the same manner as for L4D0 larvae, except that a 25 μL microinjector (Φ0.26 mm; Bolige) was used, each individual was injected with 5.0 μL of dsRNA solution, and the post-injection chilling procedure was omitted to achieve a better survival rate. The dsRNA-injected prepupae were immediately removed from the double-sided tape and placed into a small plastic cup (Φ101 × 40 mm) filled with 20 g vermiculite and 20 g water (50% water content).

### Phenotypic characterization of dsRNA-injected individuals

For *Kr-h1* knockdown (*Kr-h1*i), *Chinmo* knockdown (*Chinmo*i), and *Kr-h1* and *Chinmo* double-knockdown (*Kr-Chm*i) individuals, 2.0 μL of solution containing 2000 ng dsRNA was injected into each individual, with ds*malE* injection serving as a control. Stages of the dsRNA-treated individuals were checked 3 times a week by identifying the exuviae in the feed and measuring the head capsule width using a stereo zoom microscope (Axio Zoom.V16; ZEISS) until pre-pupation. Newly emerged prepupae (PPD0 to PPD2) were collected, weighed using an analytical balance (Entris^®^ II BCE224I-1SJP; Sartorius), and the head capsule width was recorded. Prepupae that emerged from the molt of L5 larvae were identified as precocious prepupae of *Kr-h1*i, *Chinmo*i, or *Kr-Chm*i.

For *Br-c* knockdown (*Br-c*i), 5.0 μL of solution containing 2000 ng ds*Br-c* was injected into each individual at PPD0, with ds*malE* injection serving as a control. Stages of the dsRNA-treated individuals were checked 3 times a week to confirm mobility by placing all individuals on the surface of the vermiculite and settling the cup in a dark place for 30 minutes. Immobile individuals were kept on the surface for 21 days, and then the puparium removal was performed as previously described for individuals that did not emerge as adults. These individuals were identified as *Br-c*i arrested pupae, and their morphology was recorded using a stereo zoom microscope (Axio Zoom.V16; ZEISS).

For *E93* knockdown (*E93*i), 5.0 μL of solution containing 2000 ng ds*E93* was injected into each individual at PPD7, with ds*malE* injection serving as a control. Stage identification was performed in the same manner as *Br-c* knockdown. The immobile individuals 21 days post-pupal entry were identified as E93i arrested pupae, and their morphology was recorded using a stereo zoom microscope (Axio Zoom. V16; ZEISS).

### Higher dose dsRNA-injection and analyses for *Br-c*i and *E93*i

To induce more intense morphological changes for *Br-c* knockdown (*Br-c*i), two alternative RNAi strategies were employed: (i) a single injection of 5.0 μL of solution containing 5000 ng ds*Br-c* into each individual at PPD0, and (ii) repeated injections of 3 doses of 5.0 μL solution containing 2000 ng of ds*Br-c* at PPD0, PPD3, and PPD6. The same amount and times of ds*malE* injection were conducted as a control.

To induce more intense morphological changes for *E93* knockdown (*E93*i), two alternative RNAi strategies were employed: (i) a single injection of 5.0 μL of solution containing 5000 ng ds*E93* into each individual at PPD7, and (ii) repeated injections of 3 doses of 5.0 μL solution containing 2000 ng of ds*E93* at PPD6, PPD8, and PPD10. The same amount and times of ds*malE* injection were conducted as a control.

Stage identification and phenotypic characterization were performed in the same manner as previously described for 2000 ng dsRNA injection.

### Knockdown efficiency and mRNA expression level analyses in dsRNA-injected individuals

For knockdown efficiency analysis, dsRNA-injected individuals were collected at stage-specific time points after injection. For *Kr-h1* and *Chinmo*, L4D0 larvae were injected with 2.0 μL solution containing 375, 750, 1500, or 3000 ng dsRNA and collected at L4D2 (n = 6 per group), with larvae injected with 3000 ng ds*malE* serving as the negative control (n = 6). For *Br-c*, PPD0 prepupae were injected with 5.0 μL solution containing 2000 ng dsRNA and collected at PPD2 (n = 6), with prepupae injected with 3000 ng ds*malE* used as the negative control (n = 6 per control group). For *E93*, PPD7 prepupae were injected with 5.0 μL solution containing 2000 ng dsRNA and collected at PPD9 (n = 6), with prepupae injected with 3000 ng ds*malE* serving as the negative control (n = 6).

For mRNA expression level analysis in dsRNA-injected individuals, six individuals from each treatment group were collected at the corresponding time points, with ds*malE*-treated individuals used as controls. *Kr-h1*i, *Chinmo*i, and *Kr-Chm*i individuals were collected 14 days after dsRNA injection; *Br-c*i individuals were collected 7 days after dsRNA injection; and *E93*i individuals were collected at PD3, corresponding to 6–9 days after ds*E93* injection at PPD7.

RNA extraction, cDNA synthesis, and qRT-PCR were performed as described above to quantify the relative expression levels of *Br-c*, *Chinmo*, *E93*, and *Kr-h1*, using *Actin-5C* as the reference gene. For each analysis, six biological replicates were used per group. The average relative expression level calculated using the ΔΔCt method in the ds*malE*-treated control group was normalized to 1, and all values were adjusted relative to this control for statistical analysis.

### Statistics

Two-tailed unpaired Student’s t-tests were used to compare prepupal body weight, prepupal head capsule width, and gene relative expression level between the control (ds*malE*-treated) group and the RNAi group. For the knockdown efficiency test involving control (ds*malE*-treated) and different amounts of dsRNA-treated groups, group differences were analyzed by one-way ANOVA followed by Tukey’s multiple comparisons test. To analyze fold changes in the relative expression levels of key metamorphic genes in *Kr-h1*i and *Chinmo*i individuals, group differences were analyzed by one-way ANOVA followed by Dunnett’s multiple comparisons test against the control group. All statistical analyses were performed using GraphPad Prism 10.6.1.

## Acknowledgements

We thank Yooichi Kainoh (University of Tsukuba) for providing accessibility to the tabletop SEM for photography, and Sugihiko Hoshizaki (The University of Tokyo) for his critical comments on stage identification and RNAi experiments. This work was supported by the Cabinet Office, Government of Japan, Moonshot Research and Development Program for Agriculture, Forestry and Fisheries Research and Development Program given through the Bio-oriented Technology Research Advancement Institution (JPJ009237).

## Author Contributions

Lingtao Zhang, Conceptualization, Data curation, Formal analysis, Validation, Investigation, Visualization, Methodology, Writing – original draft; Masami Shimoda, Conceptualization, Supervision, Funding acquisition, Project administration, Writing – review and editing; Chieka Minakuchi, Conceptualization, Supervision, Project administration, Methodology, Writing – review and editing.

C.M. and M.S. jointly supervised this work and serve as co-corresponding authors.

## Figure Legends

**Figure 1—figure supplement 1.**
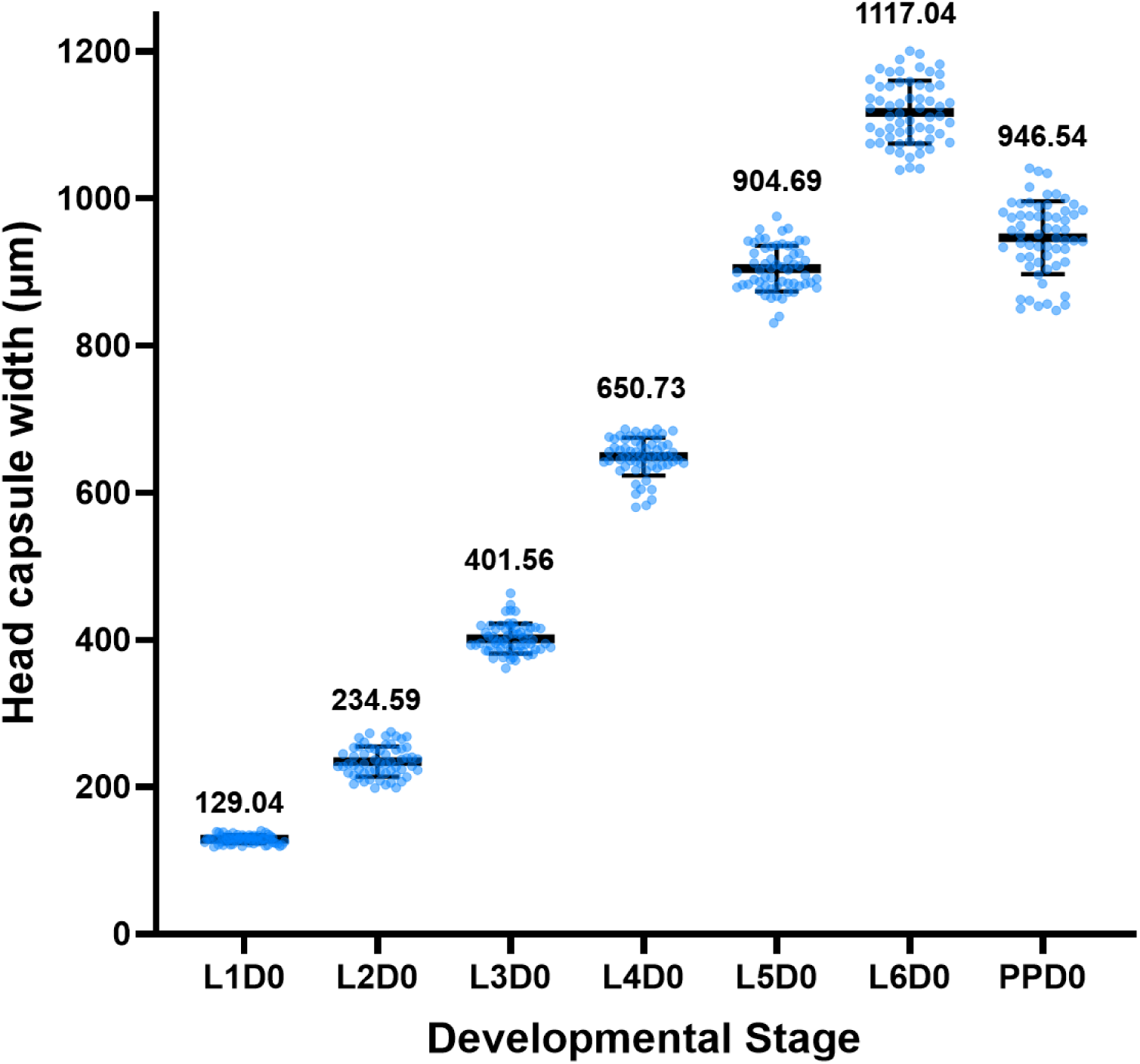
Developmental changes in head capsule width across larval and prepupal stages in *H. illucens* (mean ± SD, n = 60). **Figure supplement 1—source data 1.** Numerical values for head capsule width shown in **Figure 1—figure supplement 1.**

**Figure 1—figure supplement 2.**
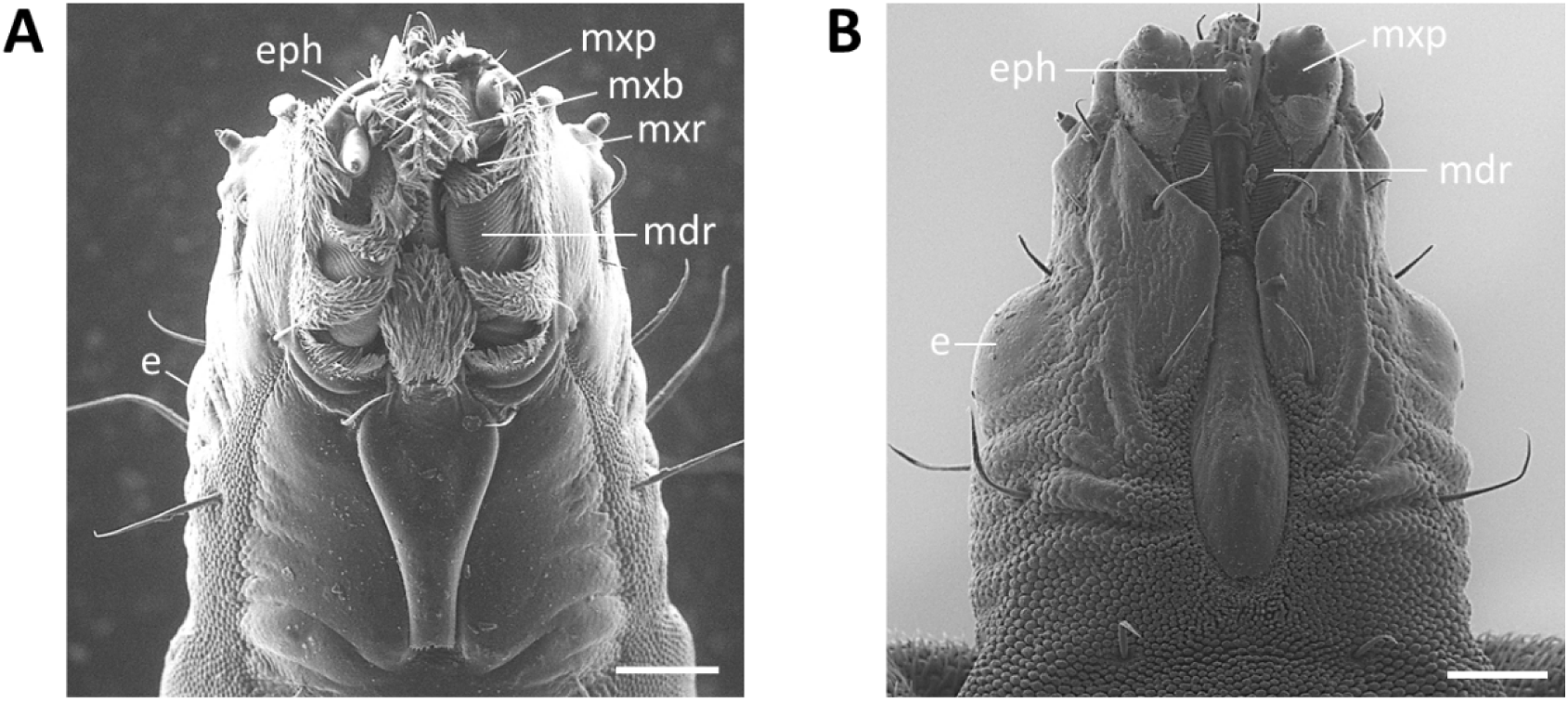
Head morphology of *H. illucens* final instar larva and prepupa. (**A**) Head morphology of the final instar larva (L6D0). (**B**) Head morphology of the prepupa (PPD0). Scale bars: 200 μm. Abbreviations: e – eye, eph – epipharynx, mdr – mandibular ridges, mxb – maxillary brush, mxp – maxillary palp, mxr maxillary ridge.

**Figure 2—figure supplement 1.**
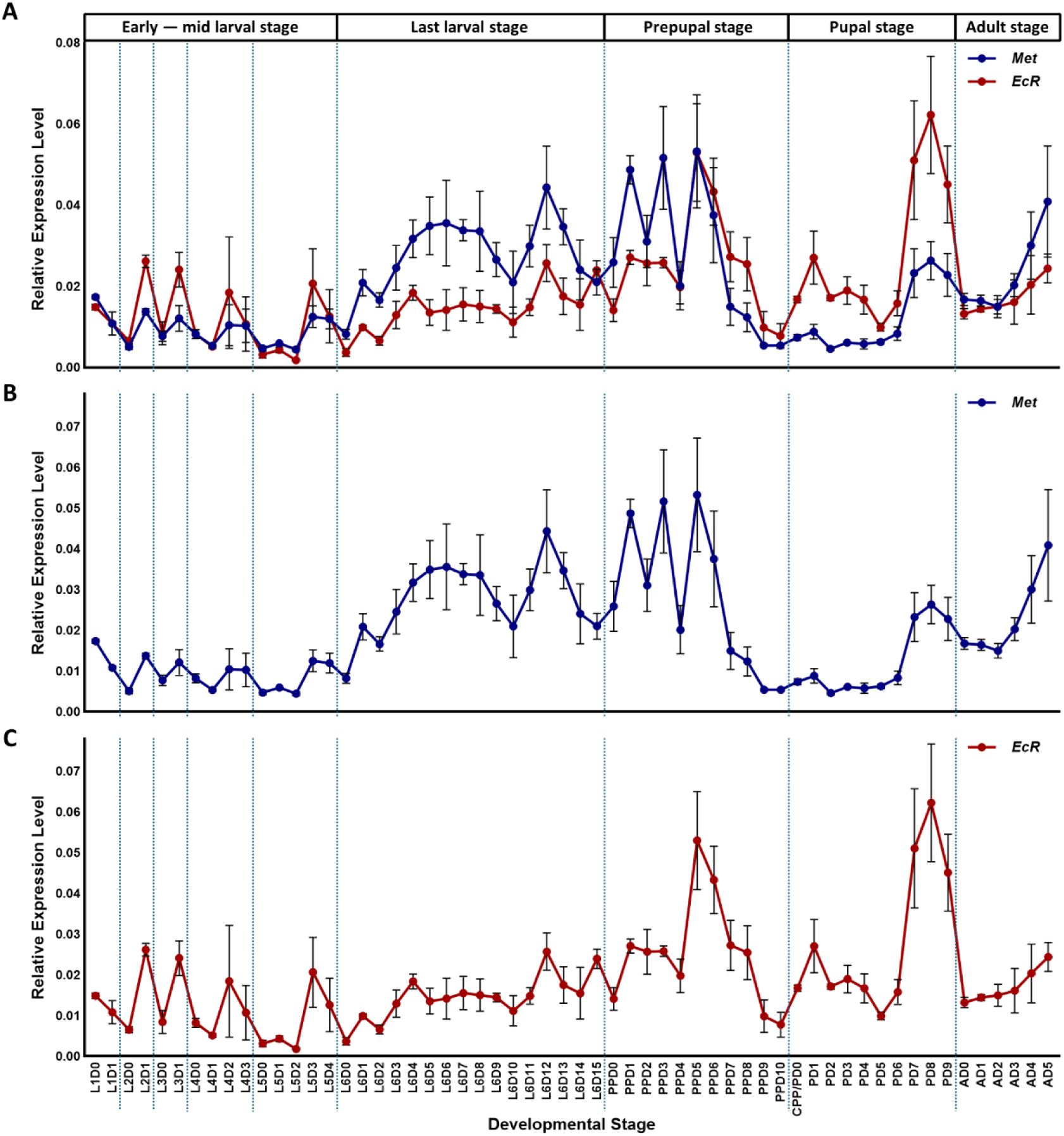
Expression profiles of genes encoding JH and ecdysone receptors. (**A**) Aligned expression profile of genes encoding JH and ecdysone receptors: *Met* and *EcR*. (**B**) Expression profile of *Met*. (**C**) Expression profile of *EcR*. Relative expression level was calculated using the 2^−ΔΔCt^ method, with *Actin-5C* as the reference gene. All data are shown in mean ± SE (n = 3). **Figure supplement 1—source data 1.** Numerical values for expression profiles shown Figure 2—figure supplement 1.

**Figure 2—figure supplement 2.**
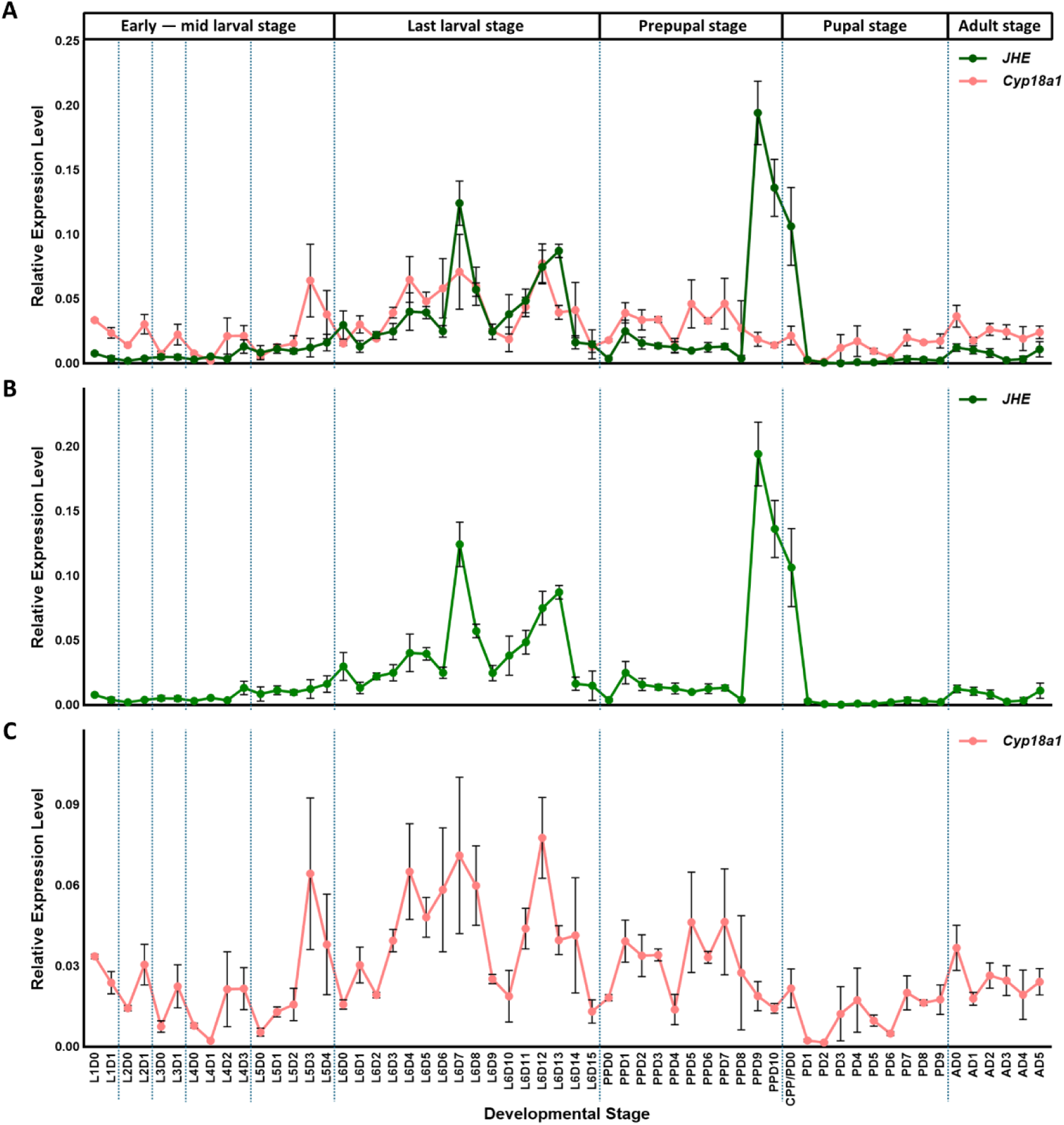
Expression profiles of genes encoding key degradation enzymes of JH and 20E. (**A**) Aligned expression profile of genes encoding key degradation enzymes of JH and 20E: *JHE* and *Cyp18a1*. (**B**) Expression profile of *JHE*. (**C**) Expression profile of *Cyp18a1*. Relative expression level was calculated using the 2^−ΔΔCt^ method, with *Actin-5C* as the reference gene. All data are shown in mean ± SE (n = 3). **Figure supplement 2—source data 1.** Numerical values for expression profiles shown **Figure 2—figure supplement 2.**

**Figure 3—figure supplement 1.**
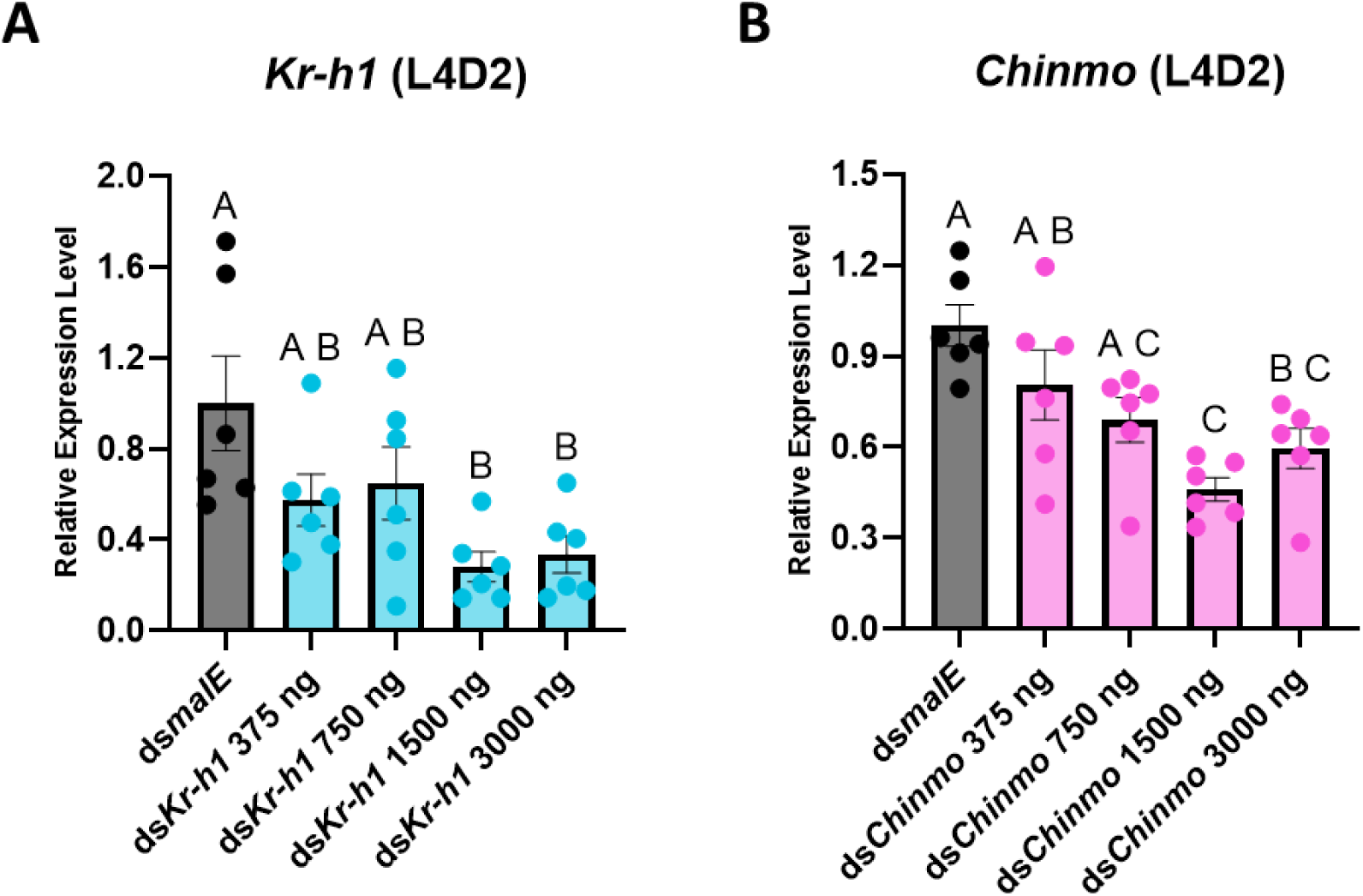
Knockdown efficiency analysis of ds*Kr-h1* and ds*Chinmo* in *H. illucens* larvae. (**A**) Knockdown efficiency analysis of ds*Kr-h1*. (**B**) Knockdown efficiency analysis of ds*Chinmo*. L4D0 larvae were injected with 2.0 μL of dsRNA solution, and collected at L4D2 for qRT-PCR analysis (mean ± SE, n = 6 for each group). Statistical analysis was performed using one-way ANOVA followed by Tukey’s multiple comparisons test. Different letters indicate significant differences among groups (*p* < 0.05).

**Figure 5—figure supplement 1.**
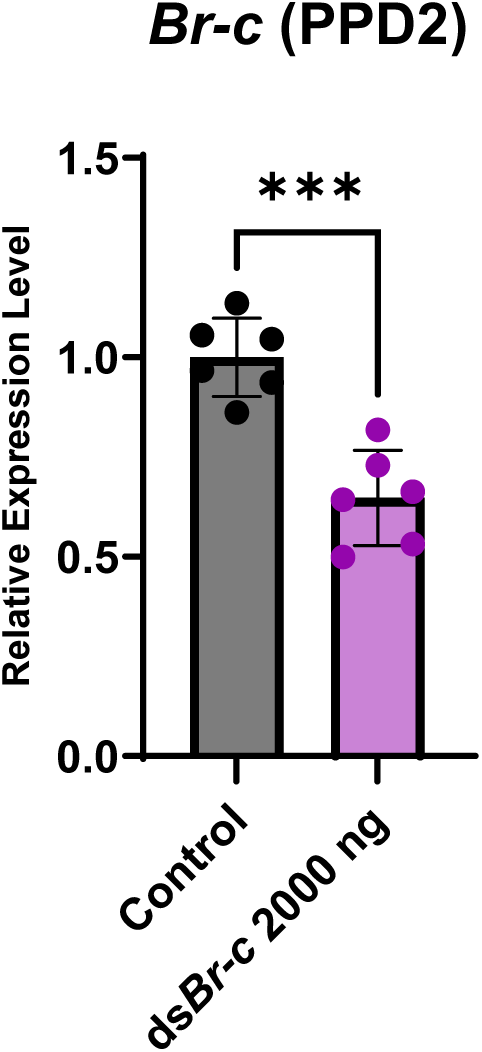
Knockdown efficiency analysis of ds*Br-c* in *H. illucens* prepupae. PPD0 prepupae were injected with 5.0 μL of dsRNA solution, and collected at PPD2 for qRT-PCR analysis (mean ± SE, n = 6 for both groups). An unpaired Student’s t-test was used to compare the two groups. Asterisks denote statistical significance: p < 0.05 (*), p < 0.01 (**), and p < 0.001 (***).

**Figure 5—figure supplement 2.**
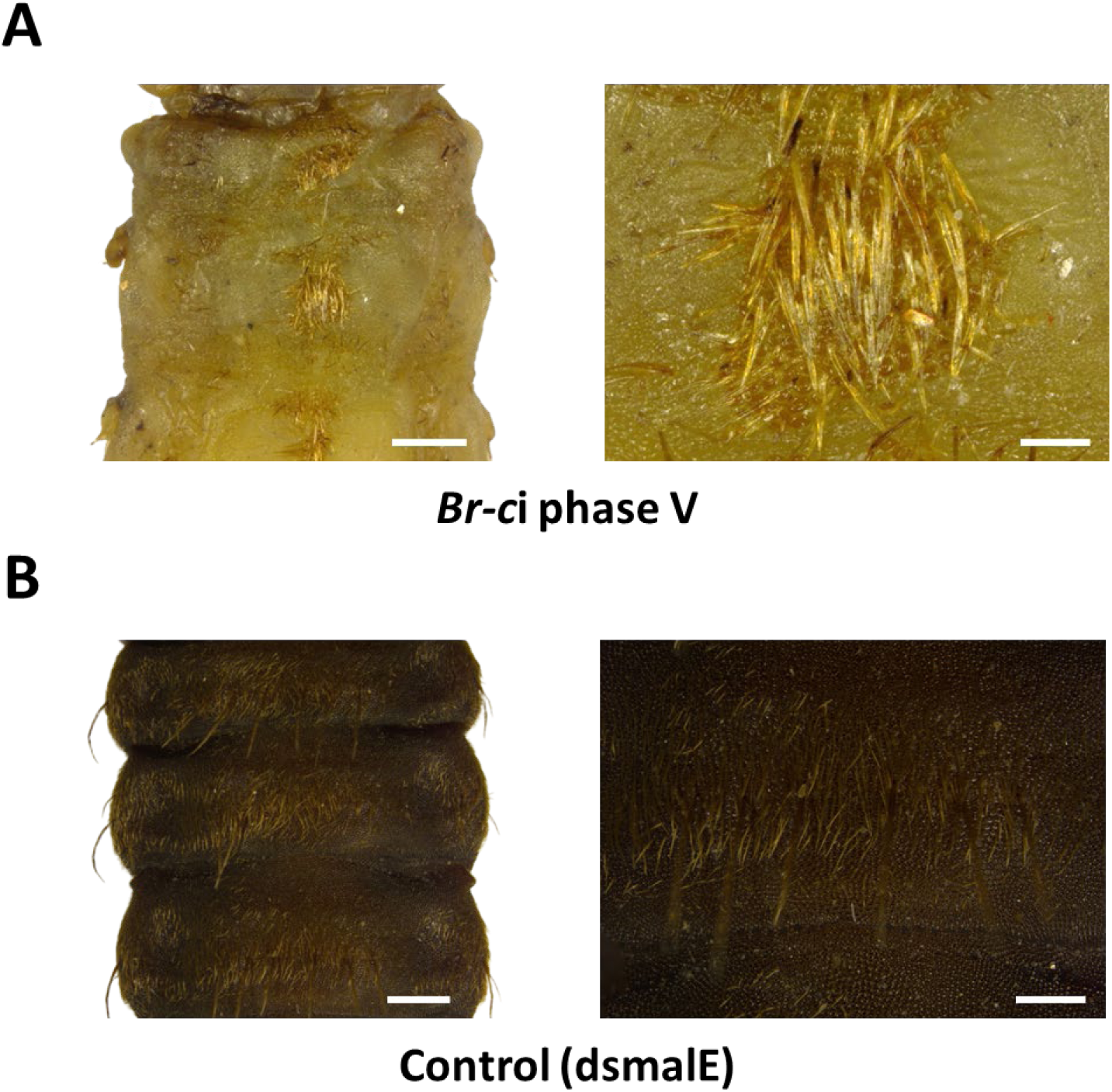
Morphological comparisons between the *Br-c*i phase V arrested pupa and the control (ds*malE* treated) prepupa’s abdomen. (**A**) ventral view of the *Br-c*i phase V individual’s 1^st^ to 3^rd^ abdominal segments (left), and the enlarged image of the abdominal setae on the *Br-c*i phase V individual’s 2^nd^ abdominal segment (right). (**B**) ventral view of the control prepupa’s 1^st^ to 3^rd^ abdominal segments, and the enlarged image of the abdominal setae on the control prepupa’s 2^nd^ abdominal segment (right). Scale bars: 500 μm for abdominal segments (left), and 200 μm for enlarged images (right).

**Figure 6—figure supplement 1.**
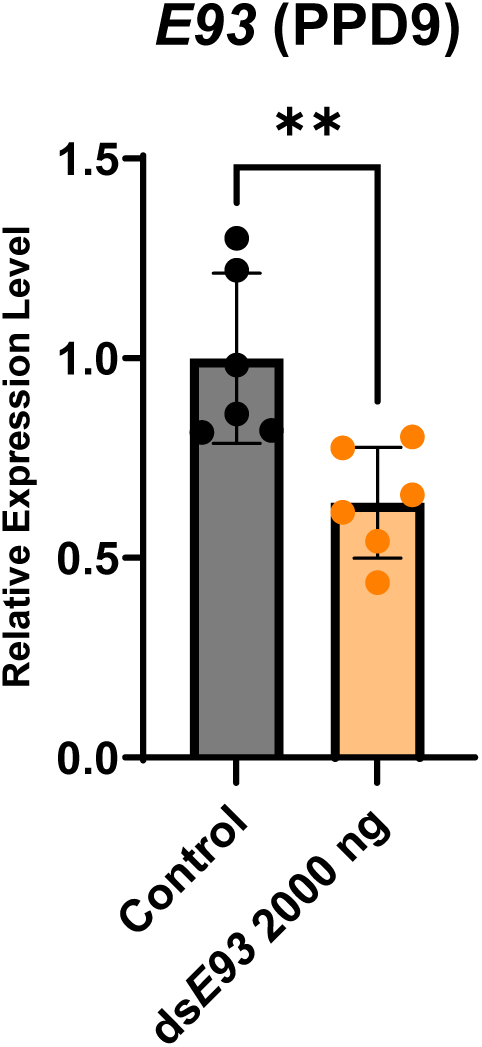
Knockdown efficiency analysis of ds*E93* in *H. illucens* prepupae. PPD7 individuals were injected with 5.0 μL of dsRNA solution, and collected at PPD9 (mean ± SE, n = 6 for both groups). An unpaired Student’s t-test was used to compare the two groups. Asterisks denote statistical significance: p < 0.05 (*), and p < 0.01 (**).

**Figure 6—figure supplement 2.**
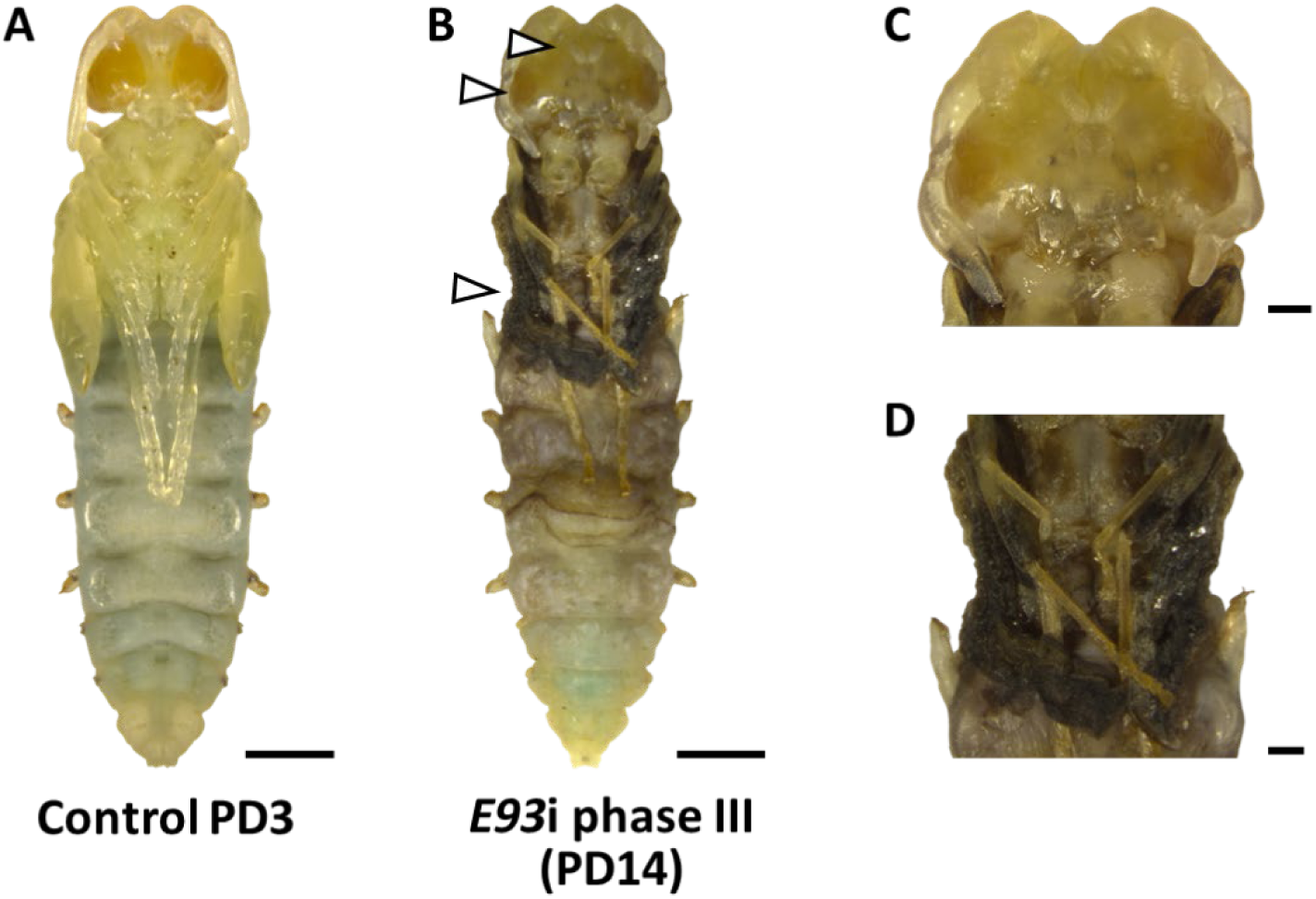
Morphological comparisons between the *E93*i phase III arrested pupa and the control (ds*malE* treated) pupa. (**A**) Ventral view of the control pupa at PD3 with amber eyes. (**B**) Ventral view of the *E93*i phase III arrested pupa at PD14: only a restricted area of the eyes was pigmented amber color; wings were partially expanded but flat, showing pigmentation in black like mature pupa; black pigmentation was also observed in legs from femora to tibiae. (**C**) Enlarged image of the head of the *E93*i phase III arrested pupa. (**D**) Enlarged image of the thorax and the 1^st^ abdominal segment of the *E93*i phase III arrested pupa. Scale bars: 2000 μm for the whole body, and 500 μm for body parts.

## Supplementary Tables

**Supplementary Table S1.**
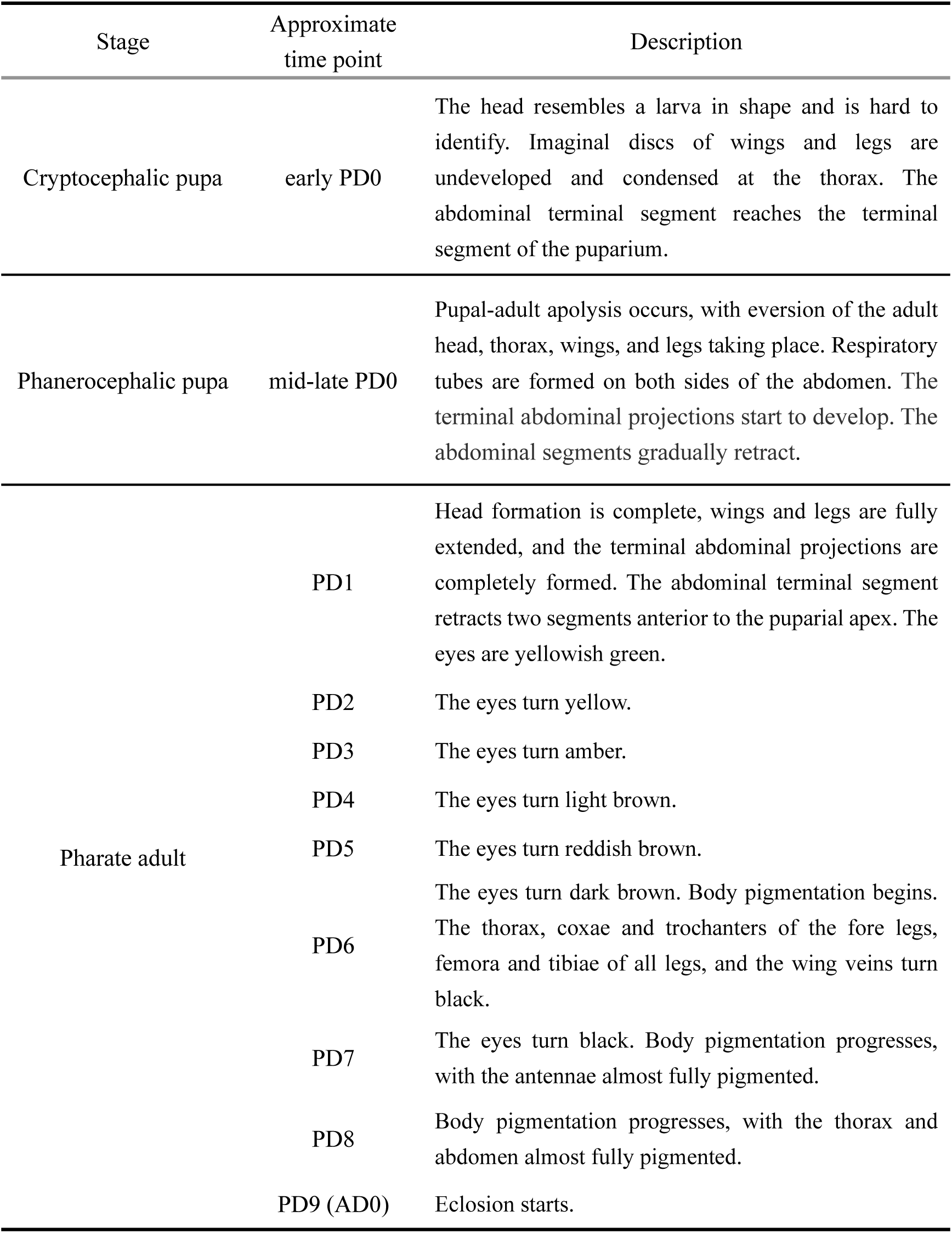
Intra-puparial developmental stages of black soldier fly.

**Supplementary Table S2.**
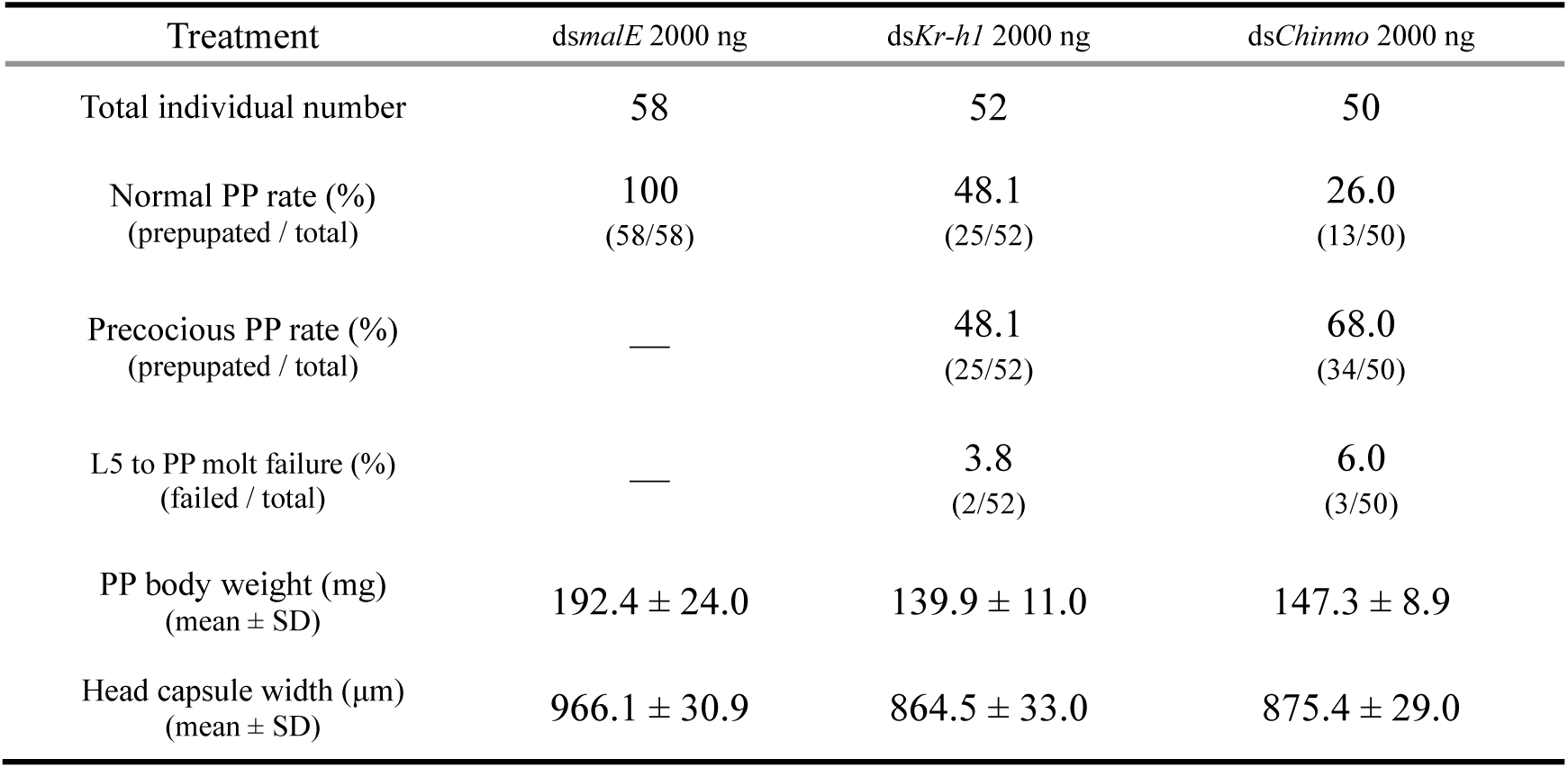
Statistical results of ds*Kr-h1* or ds*Chinmo* treatment to L4D0 larvae.

**Supplementary Table S3.**
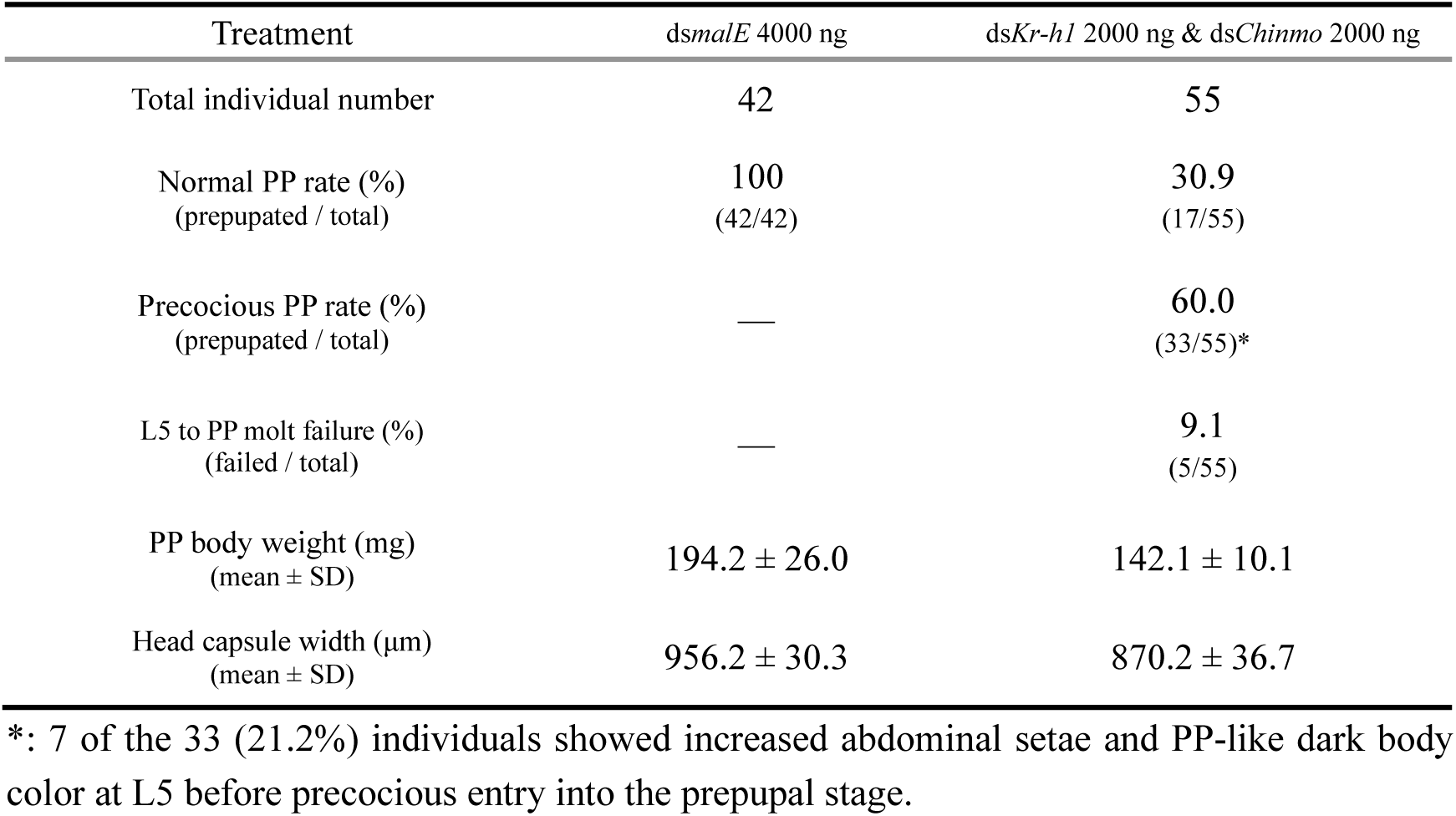
Statistical results of simultaneous ds*Kr-h1* and ds*Chinmo* treatment to L4D0 larvae.

**Supplementary Table S4.**
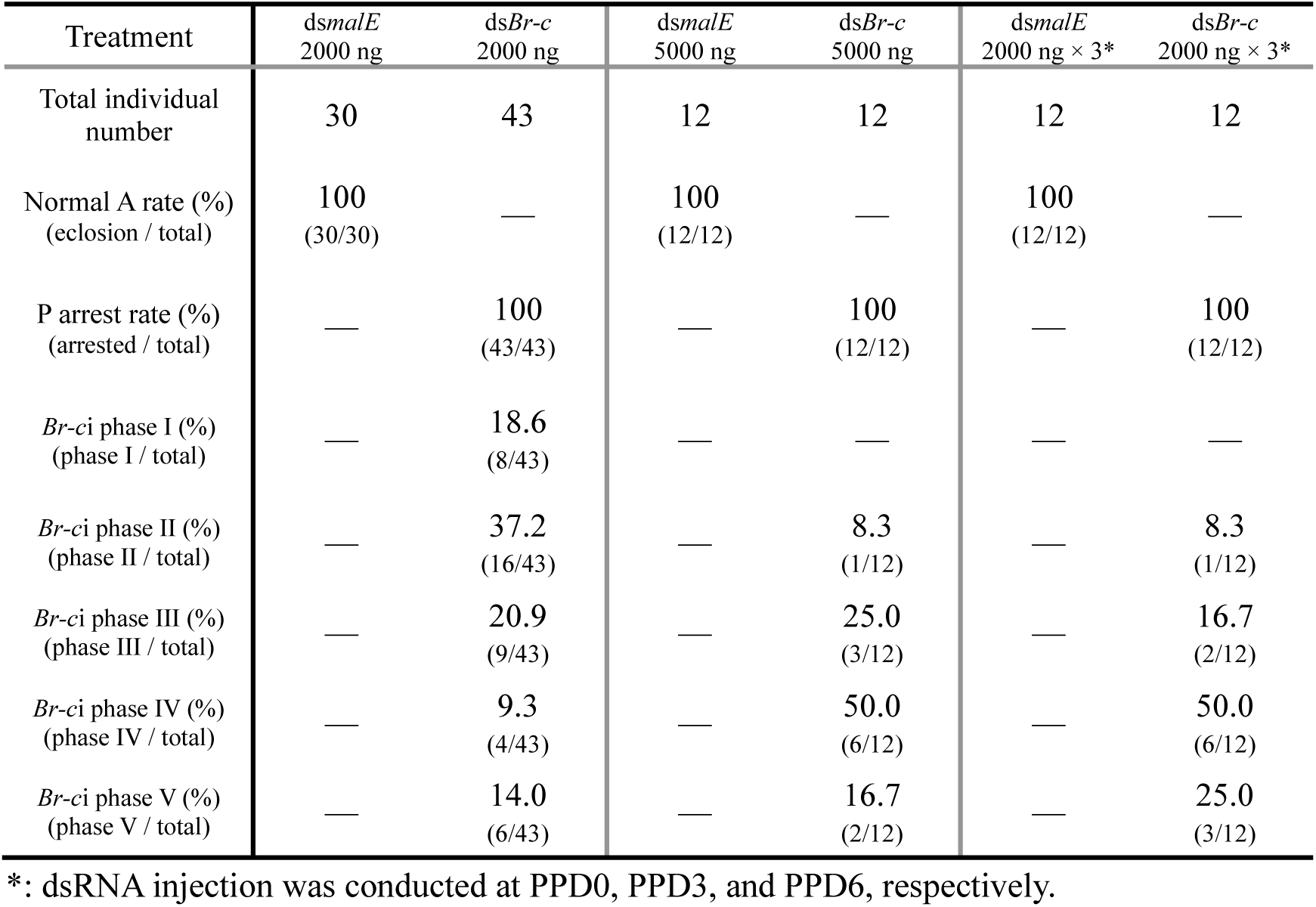
Statistical results of ds*Br-c* treatment to PPD0 (or PPD0∼6) prepupae.

**Supplementary Table S5.**
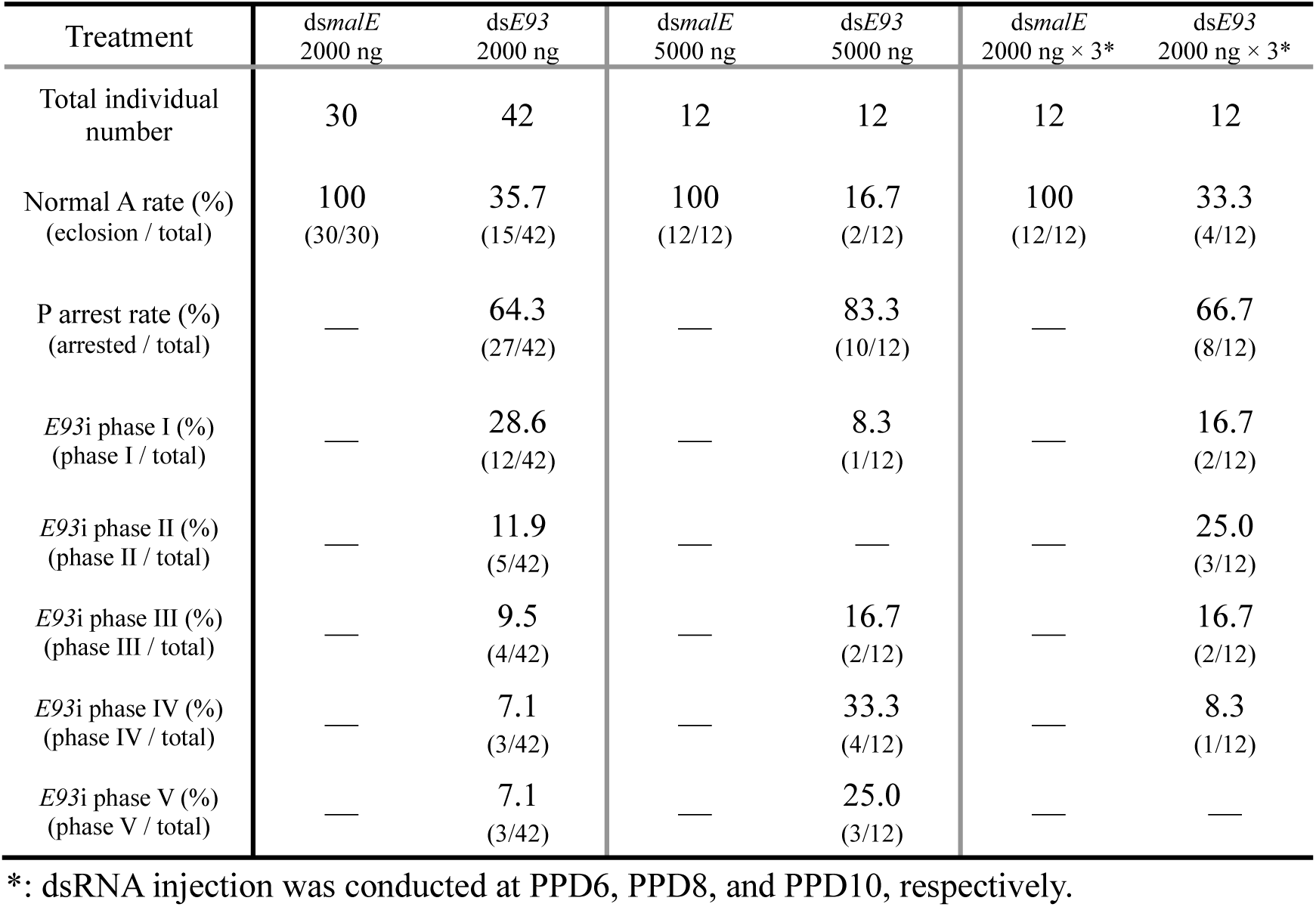
Statistical results of ds*E93* treatment to PPD7 (or PPD6∼10) prepupae.

**Supplementary Table S6.**
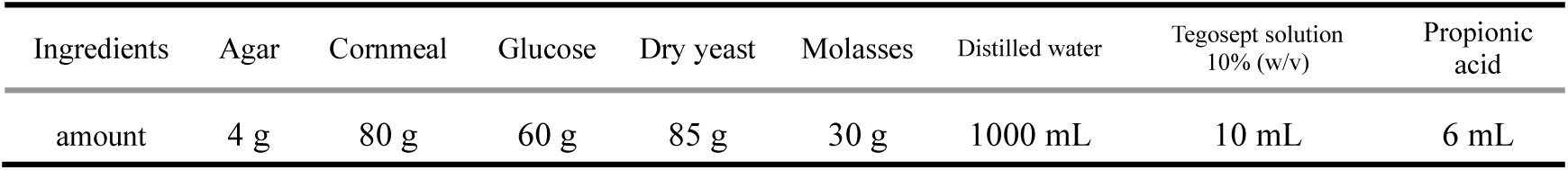
Composition of the artificial diet per liter.

**Supplementary Table S7.**
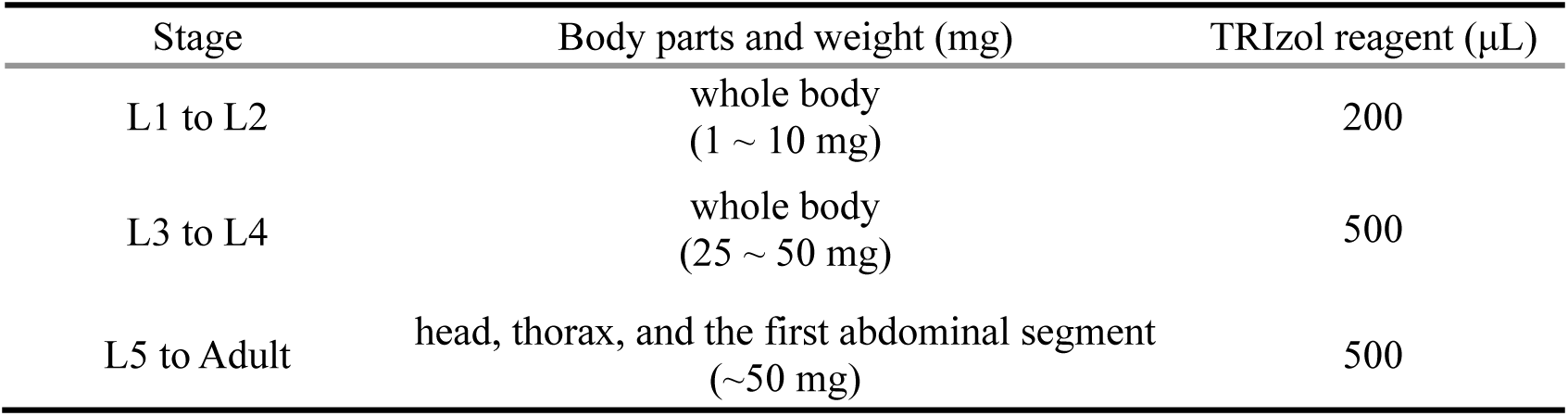
Insect body parts and the amount of TRIzol reagent used for total RNA extraction.

**Supplementary Table S8.**
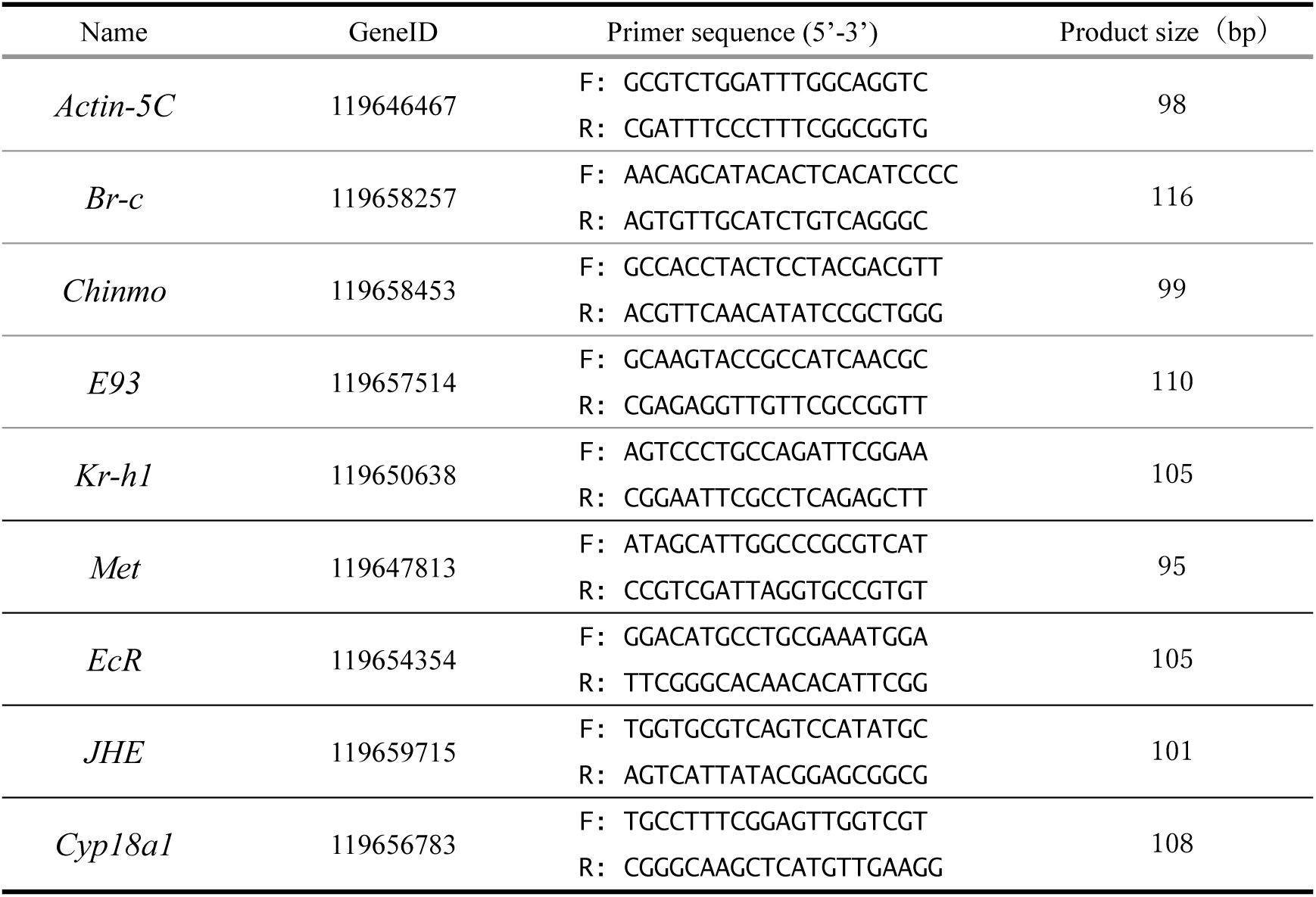
Primers used for qRT-PCR in the present study.

**Supplementary Table S9.**
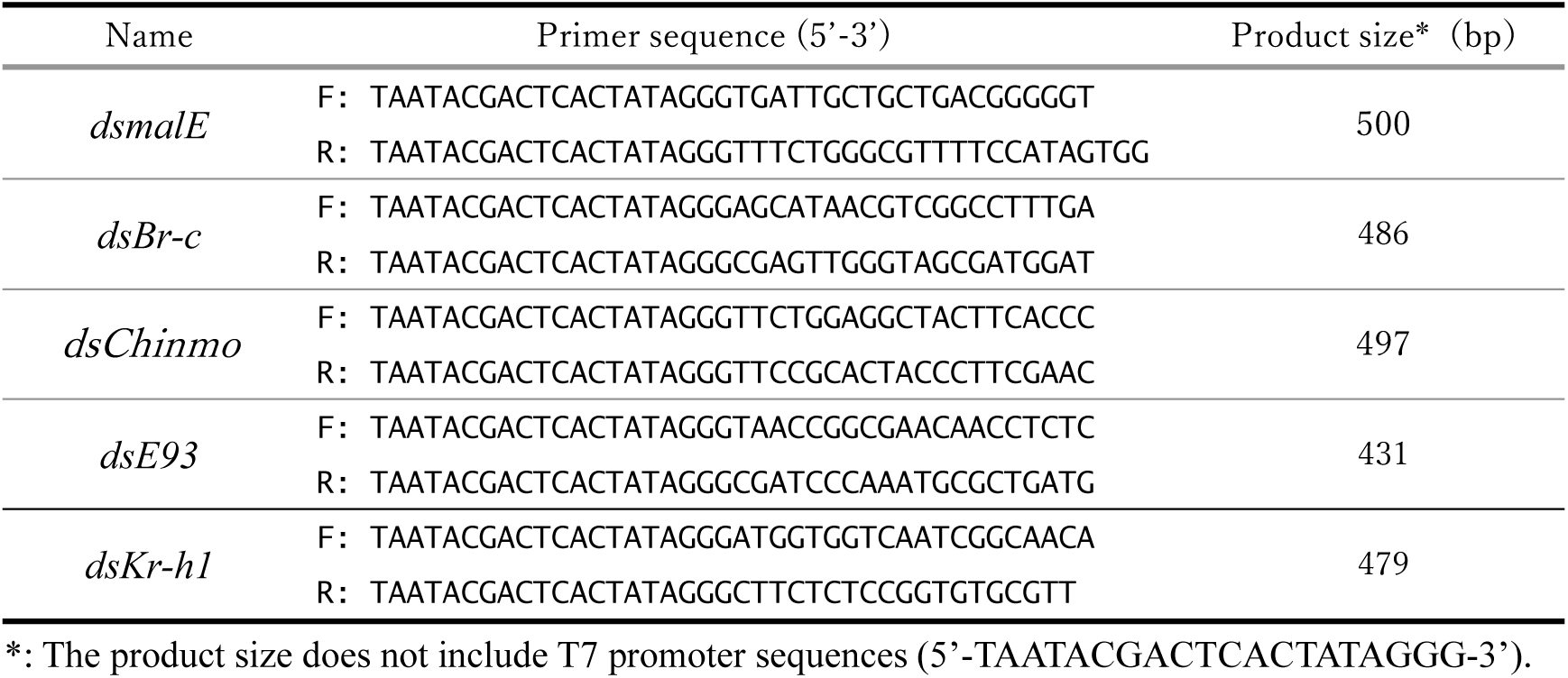
Primers used for dsRNA synthesis in the present study.

